# Hippocampal theta sweeps indicate goal direction

**DOI:** 10.1101/2025.08.21.671551

**Authors:** Changmin Yu, Zilong Ji, Jake Ormond, John O’Keefe, Neil Burgess

## Abstract

Successful spatial navigation requires rapid evaluation of potential future trajectories. Hippocampal “theta sweeps”, sequential activation of place cells within individual theta cycles, exhibit predictive dynamics within the ideal timeframe to fulfill this role. However, whether these sequences simply reflect movement-related variables or afford more cognitive goal-directed planning remains unresolved. Using data from a navigation task on the “Honeycomb” maze that allows dissociation of head-, movement- and goal-direction correlates, we found that hippocampal theta sweeps exhibit robust goal-oriented directional biases, independent of movement- or head-direction. An existing model of theta sweeps, with an additional goal-oriented directional input, reproduces these findings and predicts goal-oriented theta phase precession, which we confirm empirically. Replay events during immobility-related sharp wave/ripples are also goal-directed, and therefore more aligned with theta sweeps than experience. Our findings indicate that hippocampal theta sweeps provide a neural substrate for online goal-directed spatial planning.

## Main Text

The ability to transcend immediate sensory inputs and motor outputs to form cognitive vectors towards desired locations is a crucial aspect of successful spatial navigation and planning. Hippocampal place cells, although primarily representing the current location (*1*), exhibit remarkable sequential dynamics within the ideal timeframe for such online planning. In foraging rodents, the population of active place cells sweep forward from those with firing fields at or behind the animal to those with firing fields ahead of the animal within each cycle of the (6-12Hz) theta rhythm (*2–4*), effectively turning temporal sequences into sweeps through space (*5*) (“theta sweeps’’; Figure 1a). Recent results have suggested a cognitive role for hippocampal theta sweeps, sampling potential paths ahead of the animal, sometimes alternating to both sides of a choice point (*6–8*) or of the forward direction in open fields (*9*), and extending farther when running faster (*7*) or towards more distant (*10*) or better-learned (*11*) reward locations. However, it has been difficult to determine whether theta sweeps simply extend forwards as a consequence of movement-related dynamics, e.g., firing rate adaptation within attractor networks (*12, 13*), or whether they reflect cognition and planning, encoding “sense of direction” vectorial information towards places of interest (*14, 15*). Dissociating these possibilities is hampered by experimental limitations such as the restricted sampling of firing fields on narrow tracks, or the animal’s propensity to orient and move directly towards places of interest in open fields.

**Figure 1.**
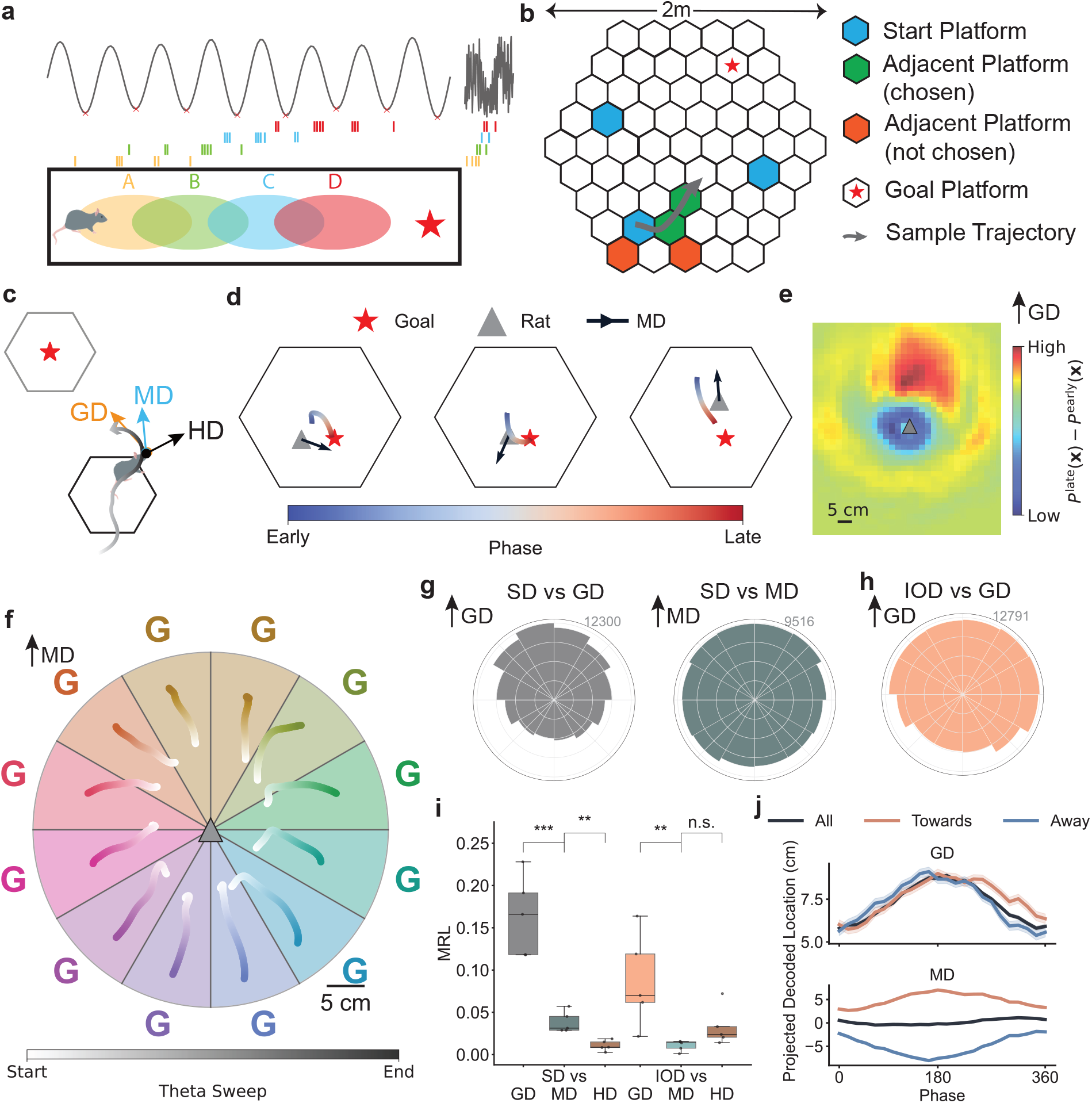
Disentangled movement and goal direction reveals strong goal-oriented direction bias in hippocampal theta sweeps. **a**. Theta sweeps during movement along linear track (left), and replay sequence during immobility (right). Red star indicates the goal. **b**. Honeycomb maze and associated transition rules (*17*). At the beginning of each trial, the rat is placed on a randomly chosen start platform (Figure S1). During each “wait-period” of ca. 10 seconds, the animal is confined to its current platform until two pseudo-randomly chosen adjacent platforms are raised, and it makes its choice by moving onto the chosen platform. Trials end upon rats reaching the goal platform. **c**. Dissociation of goal direction (GD), movement direction (MD), and head direction (HD) as a direct consequence of the behavioural paradigm on the honeycomb maze. **d**. Example theta sweeps when the rat’s MD is aligned, sideways, and opposite to the GD. **e**. Normalised difference between average decoded posteriors over the first and second halves of theta sweeps, standardised with the rat at the centre (grey triangle) and MD upwards (black arrow). **f**. Median theta sweep locations relative to the animal (grey triangle at centre) as a function of GD (coloured “G”s) relative to MD (up) for each theta cycle of the LFP. **g**. Distributions of relative sweep directions (SD) with respect to GD and MD. Histograms are constructed given all valid theta sweeps from 5 recorded rats. **h**. Same as left panel in (g), but with respect to the direction of the initial offset of the decoded theta sweeps relative to the animal’s location (IOD). **i**. Rat-specific mean Rayleigh-vector length (MRL) of the distributions in panels f and g (one-sided paired t-tests; SD wrt GD vs MD: p = 0.0004; two-sided paired t-tests; MD vs HD: p = 0.0005; IOD wrt GD vs MD: p = 0.0078, MD vs HD: p = 0.9449.). **j**. Mean decoded location (with standard error) within each theta cycle as a function of theta phase. All decoded locations are projected onto the goal direction (top) and movement direction (bottom), and separately for movement towards and away from the goal.

Here we investigate the cognitive affordance of theta sweeps by analysing place cell firing from hippocampal region CA1 of rats that have learned a goal-oriented spatial navigation task which explicitly dissociates movement and head directions from the goal direction (*16, 17*) (Figure 1b). Our analysis reveals robust goal-oriented hippocampal theta sweeps. We propose a hierarchical continuous attractor network model (*18*) with goal-directed inputs, which reproduces the empirical findings whilst making direct mechanistic predictions of goal-oriented theta phase precession (*19*), which we confirmed subsequently in the neural data. Goal-directed theta sweeps are complemented by goal-oriented replay events during immobility (*20, 21*), despite the absence of direct goalward movement trajectories. Our findings suggest that theta sweeps are a candidate neural substrate supporting flexible online cognitive planning beyond simply reflecting current movements, and likely afford offline consolidation of goal-direction.

### Hippocampal theta sweeps exhibit robust goal-oriented directional modulation

The recently developed “Honeycomb” maze enables clear dissociation of movement, head, and goal directions through task design (*16, 17*): rats navigate towards an unmarked goal through a succession of indirect intermediate platforms (Figure 1b) and often exhibit rotational scanning behaviours as they occupy these intermediate platforms, effectively dissociating movement-, head-, and goal-directions (Figure 1c; Figure S6b).

Five rats were trained to reach near-optimal performance during recording trials on the Honeycomb maze (Figure S1, see (*17*) for details). We used the population activity of all recorded CA1 place cells over valid theta cycles (105.2 ± 7.5 place cells per rat; Figure S2, Figure S3) and performed Bayesian decoding of the rat’s spatial location (*22, 23*) (see Methods) where decoded location is given by the *maximum-a-posteriori* (MAP) estimate (median decoding error given 200 ms temporal binning: 13.11 ± 2.56 cm, Figure S4). For theta sweep decoding, we used a phase-rolling temporal binning, with 60° phase window (equivalent to 14 - 28 ms), and overlapping increments of 15° (equivalent to 3.5 - 7 ms) (*24*). Due to its cyclic nature, we define a theta sweep as the most forward-extending subsequence of decoded spatial locations within a theta cycle detected given the local-field potential (LFP) (*9, 24*) (see Methods).

Visual examination of decoded theta sweeps indicates a directional bias towards the goal location, regardless of rat’s movement direction (Figure 1d; Figure S6c). Across animals, the difference between decoded positions at early and late phases of theta sweeps show stronger probability towards the goal and weaker probability closer to the animal (Figure 1e). Aggregating theta cycles as a function of goal direction relative to movement direction, a “roulette” plot clearly shows that theta sweeps indicate goal direction rather than movement direction (Figure 1f). Sweep directions exhibit strong goal-modulation and moderate movement-modulation, but no modulation from head direction (Figure 1g, i; Figure S6i, j). Residual sweep directions, after regressing out the effect of movement direction, remain significantly concentrated towards the goal direction (*25*) (shuffled test on Rayleigh concentration, p < 0.001). The direction of the initial offsets of the theta sweep locations (IOD) are also significantly aligned with the goal direction, and do not exhibit modulation by movement or head direction (Figure 1h, i; Figure S6d). Sweeps start closer to the goal when movement aligns with the goal direction (Figure S6e).

To demonstrate the differential directional-modulatory effects of goal- and movement-directions in theta sweeps, we projected decoded locations over all theta sweeps onto goal- and movement-directions (Figure 1j). Decoded locations progress along the goal direction equally when the rat is either moving towards (330° - 30°) or away (150° - 210°) from the goal, whereas the projection onto movement direction exhibits “reverse” theta sweeps when the rat is moving opposite the goal direction. Examining the full decoding posteriors yields similar findings (Figure S9g, h).

Goal-oriented directional modulation in theta sweeps are consistent for all 5 recorded rats (Figure S5), robust to decoding procedures (Figure S8), and the use of multi-unit activity rather than LFP for detecting theta rhythmicity (Figure S9).

To validate our findings in settings with more recorded place cells but less separation between head-, movement- and goal-directions (Figure S7b), we examined place cell recordings from a spatial task with alternating trials of goal-oriented navigation and random foraging in an open-field “cheeseboard’” arena (*21*) (151.75 ± 59.25 putative place cells per rat, across 4 rats, Figure S7a, c-e). We observe similar goal-modulated directional bias in both sweep direction and initial offset during goal-oriented navigation trials, and significantly weaker modulation during random foraging trials (Figure S7f-j). Movement modulation in sweep direction was significantly stronger in the open-field arena in this dataset than in the Honeycomb maze, due to the frequent alignment of movement and goal direction (Figure S7g, i; c.f. Figure 1g, i).

In summary, our findings show that, during goal-directed navigation, hippocampal theta sweeps robustly signal the direction towards a learnt goal location, with significantly weaker modulation by movement-direction, and little modulation by head-direction.

### Additional factors modulate theta sweeps

As in previous studies, we found that theta sweep length increases with the distance to the goal (*10*) (Figure 2a), potentially indicating goal-distance as well as direction. They extend farther when movement is faster (*7*) (Figure 2a), and increase with LFP theta power (Figure S6g), even after regressing out the effect of speed on theta power (*1*) (p < 0.001, see Methods). The alignment between goal- and sweep-directions becomes significantly stronger as rats engage in more active movements (stronger modulation given linear movement than head-rotations; Figure 2b. Figure S6h), and when they are farther from the goal (where it subtends a narrower range of angles from the rat) even though speeds tend to be lower (Figure 2c).

**Figure 2.**
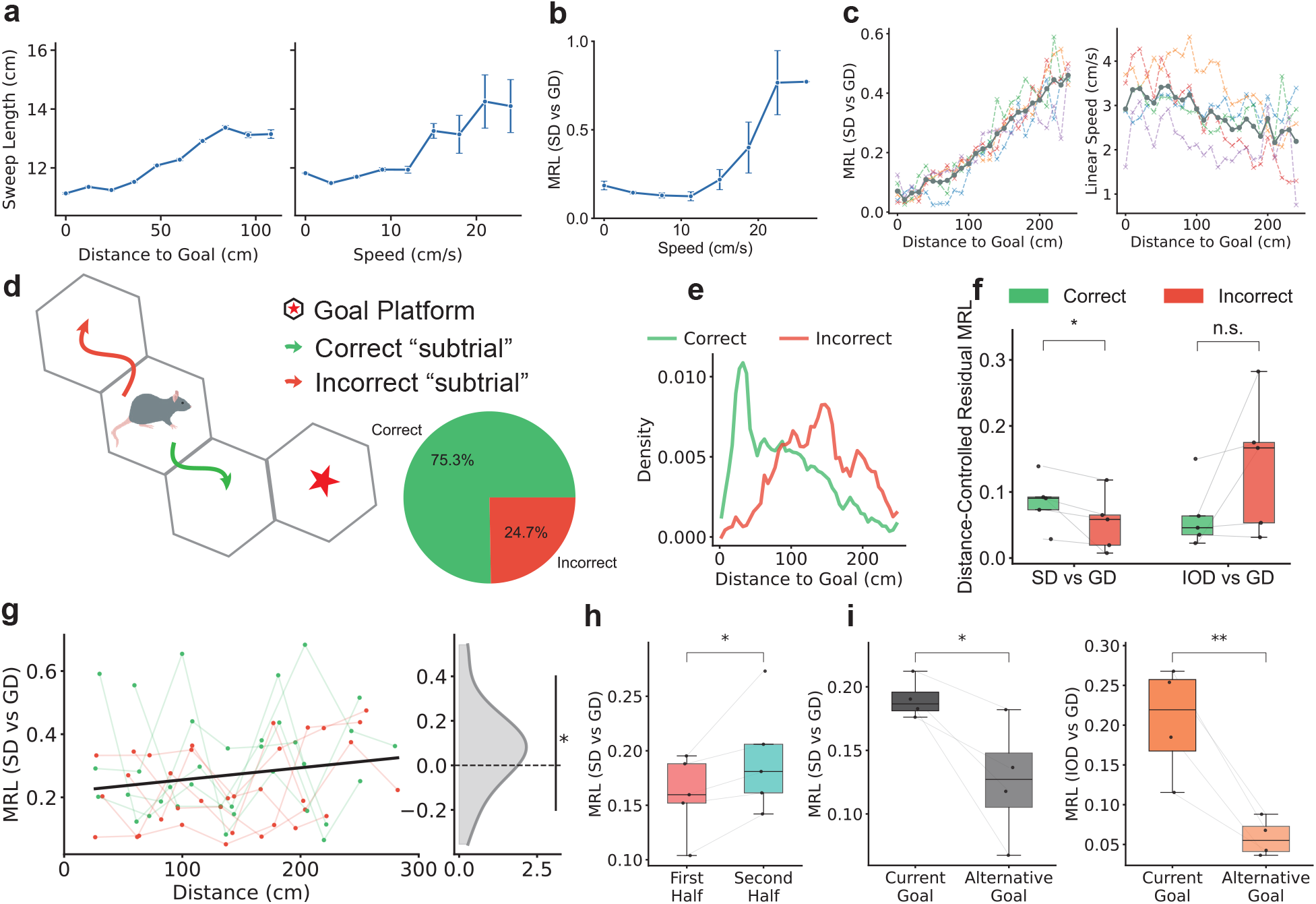
Theta sweeps are modulated by cognitively relevant behavioural correlates. **a**. Sweep lengths (mean ± standard error) as a function of distance to the goal (left) and animal’s movement speed (right). **b**. MRL of relative angles between SD and GD, as a function of animal’s movement speed. **c**. MRL of relative angles between sweep direction and goal direction (left), and movement speed (right), as a function of distance to the goal (each dashed line represents one rat, solid line represents the mean over 5 rats). **d**. Schematic illustration of correct and incorrect “choices”. Inset: proportions of correct versus incorrect choices. **e**. Probability density of rat’s distance to the goal location over correct and incorrect choices. **f**. Rat-specific MRL of distributions of residual angles after regressing out the effect of distance to the goal, for relative angles between SD and GD, and between IOD and GD (one-sided paired t-test; sweep direction: p = 0.0438; initial offset: p = 0.9251). **g**. MRL of distributions of relative angles between SD and GD, given distance-matched subsampling over correct and incorrect choices (left; black solid line represents best linear-fit between MRL and distance to the goal for all data). Residual MRL with respect to regression line over correct and incorrect choices (one-sided paired t-test: p = 0.0136). **h**. Rat-specific MRL of the circular distributions of relative angles between SD and GD, over the first and second half of all trials (one-sided paired t-test: p = 0.0269). **i**. Rat-specific MRLs for the distributions of relative angles between SD and GD (left) and between IOD and GD (right) for the current and opposing goal locations (one-sided paired t-test; SD vs GD: p = 0.0220; IOD vs GD: p = 0.0060).

Is the goalward vectorial information encoded in theta sweeps relevant for planning and capable of guiding successful navigation (*6, 8*)? We analysed “wait periods” preceding correct or incorrect choices (Figure 2d). There were no significant behavioural differences prior to correct and incorrect choices (Figure S12d-f), apart from more correct choices closer to the goal (Figure 2e). Residual sweep direction after controlling for the effect of distance to goal, either by parametrically regressing out the distance effect (Figure 2f) or non-parametrically via distance-matched subsampling (Figure 2g), exhibited significantly stronger alignment to the goal direction before correct compared to incorrect choices, while initial offsets and sweep lengths did not differ (Figure 2f; Figure S12c). Thus, the strength of goal-modulation of theta sweep direction correlates with subsequent navigational performance, complementing increases in theta sweep quality with learning of reward location on a circular track (*11*).

Goal-modulation of theta sweeps increased with experience within days, with respect to both sweep direction and initial offset (Figure 2h; Figure S10a-d), despite all rats having been extensively trained and an absence of behavioural differences over trials (Figure S1; Figure S10e-g). In goal-switching sessions, sweep directions and initial offsets were more strongly modulated by the direction towards the current goal than the alternative goal (Figure 2i; Figure S11).

Taken together, these findings reveal that theta sweeps encode cognitively relevant goalward vectorial information that correlates with subsequent navigational behaviors. The stronger goal-alignment during correct choices, progressive strengthening with experience, and alteration with goal-location shifts suggests that theta sweeps actively contribute to successful navigation. This behavioral relevance, combined with the dynamic encoding of both direction and distance to goals, positions theta sweeps as neural substrates underlying efficient and flexible online spatial planning.

### Continuous attractor network with firing rate adaptation and directional goal-modulation reproduces goal-directed theta sweeps

Forward theta sweeps in entorhinal cortical neurons, as well as their left-right alternation, can be modelled by a coupled system of continuous attractor networks (CANs) of parasubicular directional cells (DCs) and entorhinal cortical grid cells (GCs), with location-specific inputs, speed-regulated theta modulation, and firing rate adaptation mechanisms (*12, 13, 18, 26*). To model theta sweeps in hippocampal place cells (PCs), we propose a hierarchical extension of the model (*18*), incorporating a 2D attractor PC-network, receiving afferent inputs from the GC-network (Figure 3a). To induce goal-modulation, we simulate a top-down goalward allocentric directional input to the DC-ring-attractor (see Methods), to bias the DC-network activity bump toward the goal direction during each theta cycle, which subsequently activates the downstream conjunctive GCs with shared preferred internal directions with activated DCs (*27*). The conjunctive GC-network drives the activity bump in the GC-attractor towards the goal-direction. Finally, the GC activity is projected onto the PC-network through one-to-one linear projection, biasing sequential activation of PCs (Figure 3d) along the goal direction under theta modulation.

**Figure 3.**
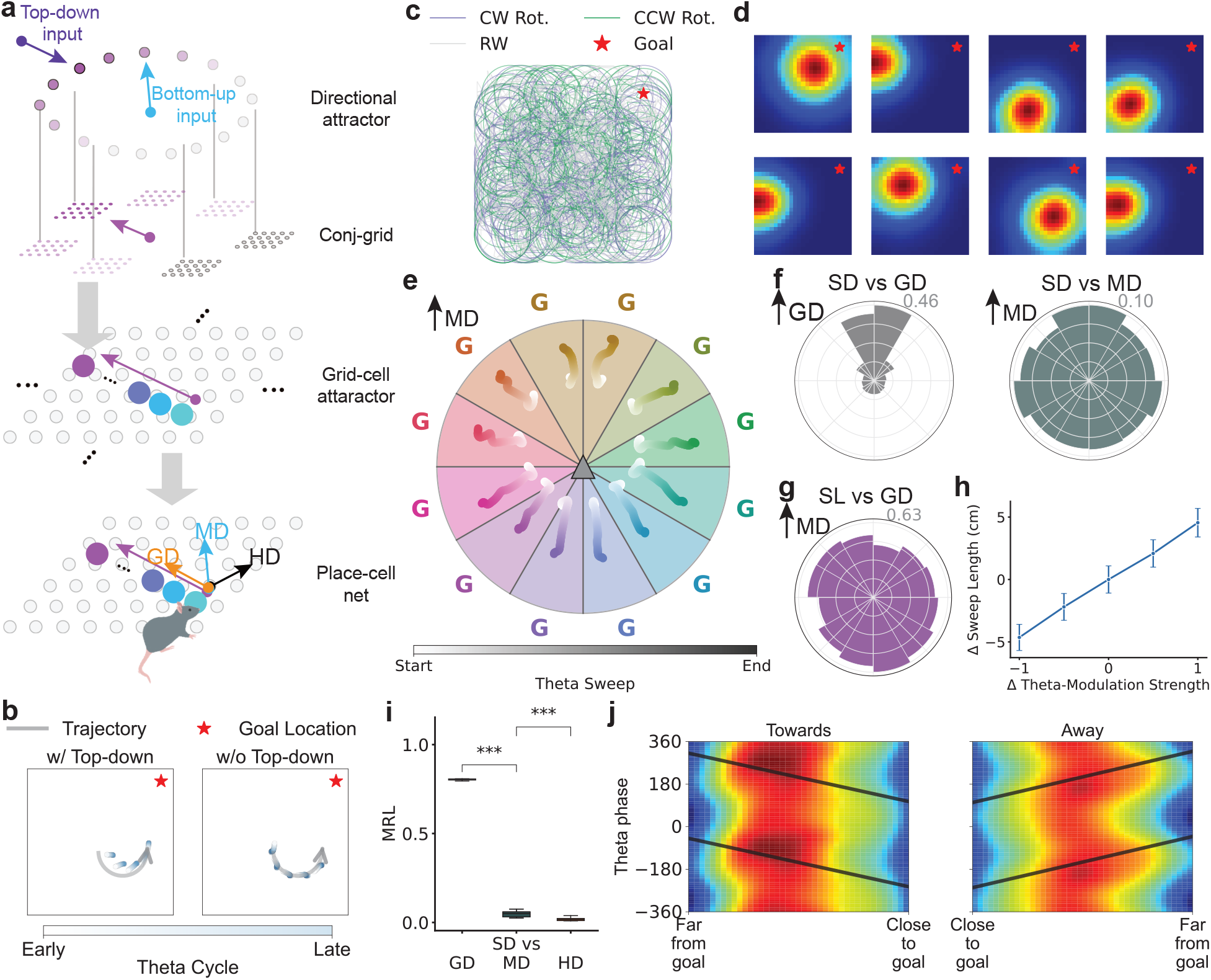
A continuous attractor network with firing rate adaptation and goal-direction input captures goal-oriented theta sweeps. **a**. Schematic illustration of the hierarchical mechanistic model. The highest-level directional cells (DC) ring-attractor network receives input of animal’s movement direction, and a top-down allocentric directional input towards the goal location. The DC attractor projects directional input with goal-oriented bias to the conjunctive grid cells (*27*), and subsequently drives phase-shift in the activity bump in the downstream grid-cell (GC) attractor network along the goal direction. The GC attractor projects linearly onto the hippocampal place cells (PC), driving population PC activity bump towards the goal direction. Orange, cyan, and black arrows indicate the goal-, movement- and head-directions, respectively. **b**. Simulated theta sweeps (forward-extending subsequence) with and without top-down goal-oriented directional input. **c**. Sample trajectory used in simulation, consists of random walks (RW) and significant periods of rotational “scanning” movements (both CW and CCW). **d**. Example spatial tuning of simulated place cells. **e**. Median simulated theta sweep locations as a function of GD (coloured “G”s) relative to MD. **f**. Circular histograms of SD relative to GD (left) and MD (right). **g**. Median sweep lengths as a function of GD relative to MD given model simulation (left) and in real data (right). **h**. Mean sweep length (± 1 s.e.) as a function of increasing theta-modulation strengths. **i**. MRL of distributions in panel g (averaged over 10 random seeds; one-sided one-sample t-test; SD vs GD: p = 2.3541 × 10^−38^; SD vs HD: p = 2.2197 × 10^−35^). **j**. Theta phase as a function of normalised distance to the goal (ND) during continuous movement periods towards (left) and away from (right) the goal. Black lines represent (warped) best-fit circular-linear regression line between ND and theta phase.

In our simulations, goal-oriented top-down directional inputs to the DC-network are necessary for driving the generation of goal-directed theta sweeps (Figure 3b).

Given simulated trajectories resembling those of rats in the honeycomb maze (interleaved random walks and rotational scanning periods, resulting in frequent non-goalward movements; Figure 3c; Figure S6a; Figure S13a; see Methods), the model successfully reproduces our key findings: spatial decoding of simulated theta sweeps, and their initial offsets, exhibit strong goal-oriented directional bias, and little effect of movement- or head-directions (Figure 3f, i). By contrast, simulated theta sweeps revert to forward and left-right alternation around movement direction without the top-down goal-directed input (Figure S13b-e). A direct consequence of instantiating theta-modulated DCs and PCs is the positive scaling of simulated sweep lengths with theta power (modelled by increased theta modulation strengths; Figure 3h), which is consistent with the neural data (Figure S6g). Moreover, the model makes the non-trivial prediction of longer theta sweeps during movements away from the goal (Figure 3g), which we also confirmed empirically in the neural data (Figure S13f). Mechanistically, this is due to the joint effect of concurrent movement of the animal during the course of goal-directed sweeps, and reduced firing rate adaptation due to misalignment of sweep- and movement-directions (compared to goalward movements).

### Goal-oriented theta phase coding

In rodents running on linear tracks or open fields, place cell firing exhibits “theta phase precession”, in which place cells fire at late phases of the theta rhythm when their firing field is ahead of the animal and at successively earlier phases as the animal moves through the field (*19*), consistent with a higher intrinsic firing frequency than the LFP theta (*28, 29*). Thus, the appearance of theta sweeps in all but the first run in a new environment (*30*), could be a direct population-level consequence of theta phase precession (*2, 3*). However, the goal-directed sweeps here require a population-level description, as above, and support an additional population-level cause of phase precession, as per the models cited above. In this case, the goal-directed theta sweeps predict theta phase *precession* when the animal is moving towards the goal, and theta phase *procession* when the animal moves away from the goal (Figure 3j). We confirmed this prediction empirically, finding strong directionally dependent phase precession and procession with respect to distance traveled along the movement direction either towards or away from the goal, respectively (Figure 4a-c).

**Figure 4.**
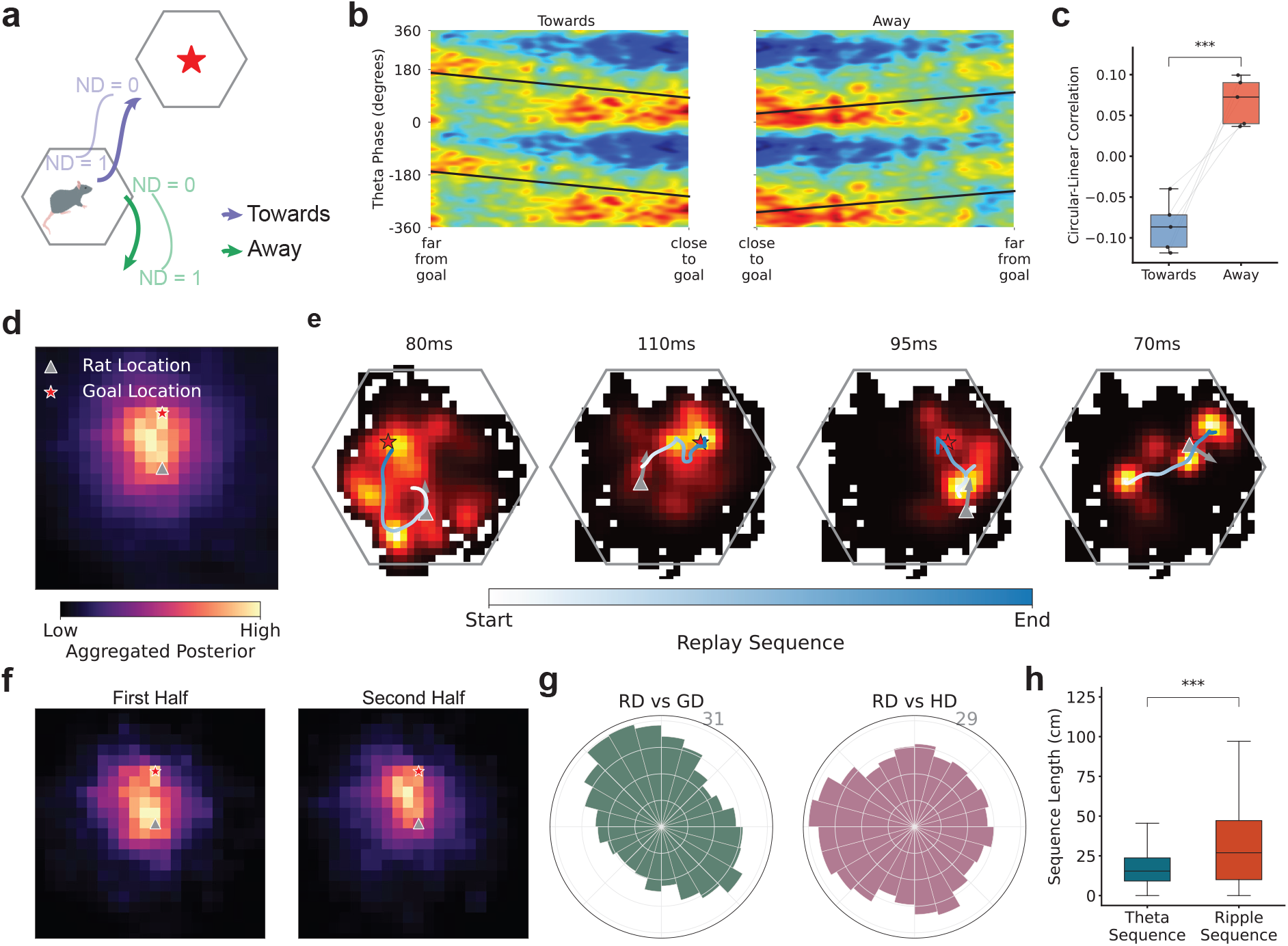
Goal-oriented sequential dynamics beyond theta sweeps. **a**. Example trajectories towards and away from the goal (star platform), illustrating the normalised distance to the goal (ND, see Methods). **b**. Theta phase as a function of ND for towards (left) and away from (right) trajectories, aggregated over all recorded putative place cells across all rats (circular-linear correlation between theta phase and signed ND; towards: r = -0.0519, p < 0.001; away: r = 0.0142, p = 0.0144; Rayleigh circular-circular test (*25)*). **c**. Rat-specific circular-linear correlation coefficients between theta phase and signed ND to the goal, for towards and away trajectories (two-sided paired t-test: p = 0.0002). **d**. Average decoded posterior probability of all replay events, standardised with respect to the rat location and direction to goal, and scaled according to the distance to the goal location. **e**. Example replay trajectories (from light to dark blue) with summed posterior probability. Gray arrow head denotes rat’s head direction. Time at the top indicates duration of the replay events. **f**. Average decoded posteriors (standardised and scaled) during the first (left) and second (right) half over all candidate ripple events. **g**. Distributions of direction of replay sequences (RD; see Methods), relative to goal- (left) and head- (right) direction, upwards in each plot. **h**. Distributions of lengths of theta sweeps and replay sequences (Mann-Whitney U-test: U = 56414443.0, p < 0.001, effect size: r = 0.2958, see Methods).

### Goal-oriented hippocampal replay sequences

Hippocampal “replay” is another form of sequential neural representation in place cells (*20, 21*), occurring during immobility (either awake or asleep) and associated with sharp wave/ripples (*1*) instead of theta rhythmicity (*31*). Replay events in open-field arenas exhibit a goal-directed bias (*21*), require LTP during encoding (*30*), and contribute to memory consolidation (*32, 33*). Replay sequences are generally thought to reflect recent experience (*34*), including those leading to reward consumption (*35*). However, the fact that replay is contingent on theta rhythmicity during encoding (*36, 37*) and includes locations that have not been physically visited (*38*), combined with the way theta sweeps afford plasticity between place cells with adjacent firing fields (*3, 19*) suggests that replay might reflect spatial trajectories composed of theta sweeps rather than experience (*11, 39*).

We examined periods of awake immobility on the Honeycomb maze (speed < 2 cm/s, z-scored multi-unit activity > 3) and identified candidate replay events (see Methods). For aggregated analysis, we rotate and scale each decoded replay event such that the goal location is always at a fixed distance north of the rat’s location. We observe over-representation of decoded posterior probability towards the goal location and moving closer from the first to second half of each event (Figure 4d, f). Visual examination of individual replay trajectories corroborates the population analysis (Figure 4e). Replay trajectories preferentially align with the goal direction (Figure 4k), whereas experienced movement- and head-directions do not over-sample the goal direction (Figure S6b). Replay sequences are significantly longer than theta-sequences (Figure 4k), suggesting that replay trajectories reflect the composition of multiple theta sweeps, consistent with suggestions that replay trajectories reflect LTP in CA3 recurrent connections during theta-sweeps (*11, 39*).

## Discussion

Our results reveal robust goal-oriented directional modulation in hippocampal theta sweeps, stronger during correct choices, suggesting that they reflect a location-independent sense of direction towards the goal, beyond simply sampling possible future movement directions (*4, 6, 8, 40*). Importantly, variation in goal-modulations is reflected in subsequent navigational performance. Hence, theta sweeps may neurally implement a key element of the cognitive map theory (*1, 41*), encoding the goalward “cognitive vector” for planning. These cognitive vectors transcend immediate sensory experience and encode trajectories towards desired future states irrespective of movement constraints. Complementing fan-tail representations of preferred goal directions by place field directional firing rates (*17*) or other goal-oriented modulations (*42, 43*), theta sweeps potentially support both flexible planning strategies and goal-related plasticity within the neuronal assembly through temporal coding schemes (*11, 39*).

Building upon an existing continuous attractor network model that captures forward and left-right alternating theta sweeps in parasubicular-entorhinal circuitry (*18*), we propose a hierarchical extension to place cells, augmented with goal-directed top-down input to the directional attractor which successfully reproduces the characteristics of goal-directed theta sequences. We confirmed phase-coding predictions from this model in the neural data, supporting the validity of this type of model (*12, 13, 18, 26*). A key assumption of the model is the presence of a goal-directed input to the theta-modulated directional cells, which switches theta sweeps from alternating forward sweeps to goal-directed sequences. The source of this input is unknown, but a promising candidate would be retrosplenial cortex (RSC), which encodes prospective goalward trajectories in both allocentric and egocentric reference frames (*44, 45*). Such integrated self-motion and goal-information might originate further upstream given projections from posterior parietal cortical neurons (*46, 47*). Albeit being essential to the proposed model, the top-down goal-directed input through subicular directional cells is not necessarily the only source conveying goal-information that drives goal-directed hippocampal theta sweeps. Alternatively, medial prefrontal cortex could project information about the action-plan/goal to the hippocampus via the thalamic nucleus reuniens (*48, 49*).

Our observation of goal-directed replay sequences during rest on the Honeycomb maze, in which there are no direct trajectories, further supports the idea that replays are composed of theta sweeps from online experience (*30, 36, 37, 39*), potentially reflecting their affordance of plasticity across place cell assemblies (*3, 19, 50*). Thus offline replay events may consolidate desired trajectories encoded by theta sweeps during active locomotion into long-term spatial knowledge, potentially supporting subsequent planning (*38*).

The conservation of theta rhythms and place cell-like representations across mammals suggests that goal-directed sequential processing represents a fundamental computational principle that emerged early in mammalian evolution (*51–53*). The goal-directed capacity of theta sweeps established here may represent predictive representations of task-relevant locations at later theta phases (*54–56*). The precise role of theta sweeps in supporting spatial planning could be causally assessed via interventions targeting specific theta phases or theta-related sequential dynamics in the entorhinal-hippocampal circuitry (*37*). The results suggest a plausible mechanism for the goal-direction function of the hippocampal cognitive map (*1*).

## Acknowledgments

We thank Brad Pfeiffer and David Foster for providing access to the neural data used in the open-field analysis, and Máté Lengyel for useful discussions.

## Author contributions

Conceptualisation: CY, JO, JOK, NB

Neural data analysis: CY, with input from JO and NB

Mechanistic modelling and simulations: ZJ, CY, with input from NB

Supervision: JOK, NB

Writing – original draft: CY, NB

Writing – review & editing: CY, ZJ, JOK, NB

## Supplementary Materials

### Materials and Methods

#### Behaviour and Electrophysiological Data

##### Honeycomb maze

Here we briefly summarise the experimental and data collection procedures used in our main analysis, and we refer readers to (*17*) for detailed description. Five male Lister hooded rats, aged between 9 to 12 months at time of electrophysiological recordings, are trained to navigate to a session-specific fixed reward location from different initial location in the Honeycomb maze, up to the point when rats were consuming food on the reward platform without hesitation (after 1 or 2 days; Figure S1a-e). The Honeycomb maze consists of 61 tessellated hexagonal platforms in an overall hexagonal configuration (200 cm in width). Each platform can be independently raised and lowered, and the raised position is ∼30 cm above the lowered position. After the initial platform was selected and raised, the animal was manually placed on the platform. Two randomly selected adjacent platforms are raised such that at least one of the platforms provides a position closer to the goal location than the current platform. After the animal has made its choice (indicated by continuous presence on the platform for 5 seconds), the remaining two platforms are lowered. In some choices, the two choice platforms are of equal distance to the goal, which we preclude from the analysis associated with correct and incorrect choices (Figure 2d-g, Figure S12). For the single-goal sessions used in main analysis, each animal was recorded over two sessions on the Honeycomb maze, with 20 trials for rats 2 and 3, and 26 trials for rats 1, 4, 5. Each trial last between 15.63s and 614.84s (164.10 ± 107.02s, averaged over all 118 trials). For goal-switching session, all Rats are firstly recorded over 13 trials to goal 1 (same goal as in the single-goal sessions), followed by interleaved “easy” trials guiding animals to the alternative goal location and “no-reward” trials that guide the animal back to the original goal location but without reward. Rats 1-4 have been successfully trained to navigate towards new goal locations (Rat 2 requires training over 6 days), and is subsequently recorded for 13 trials post-switching within the same session. Rat 5 persisted in going towards the original goal after extended training, and is hence excluded from all associated analyses (Figure S11).

Neural recordings were obtained via an electrode array with 32 gold-plated tetrodes in the dorsal CA1 region of hippocampus, with recording frequency of 20 kHz for Rat 1, and 30 kHz for Rats 2-5. Number of neurons recorded per rat ({# putative place cells}/{# all recorded neurons}): Rat 1: 98/110; Rat 2: 112/132; Rat 3: 117/126; Rat 4: 114/128; Rat 5: 95/107. Behavioural recordings are based on two infrared LEDs positioned on top of the animals’ implant (one in the front and one behind, along the rostral-caudal axis), sampled at 25 Hz. Tracking of the LEDs was performed with DeepLabCut (*57*). Vectors connecting the two LEDs determine the animal’s location (midpoint) and head direction.

#### Open-field cheeseboard arena

We briefly summarise the experimental and data collection procedures in the open-field cheeseboard arena (Figure S7), and we refer readers to (*21*) for detailed description. Four male adult Long-Evans rats, aged 10-20 weeks old, were trained on a foraging task in a 2m × 2m open-field arena, with 36 identical, evenly spaced reward wells over a 6 × 6 grid. All rats receive training (in addition to pre-training on a linear track in a separate room) to perform an alternating homing-foraging task, where a session-specific fixed home well is initially filled at the start of the session. Once the rat discovered and consumed the reward at the home well (homing trial), the foraging trial starts immediately and a randomly chosen well is then filled for the rat to discover and consume. All 4 rats are recorded over one 30-minute sessions per day, over two days.

Upon reaching a performance threshold on the task, neural recordings were obtained via a microdrive array with 40 gold-plated tetrodes implanted in the CA1 region of dorsal hippocampus, with recording frequency of 32,556 Hz. Number of neurons recorded per rat (session 1 | session 2): Rat 1: 204/213 | 253/263; Rat 2: 174/181 | 153/164; Rat 3: 79/80 | 79/80; Rat 4: 94/97 | 178/186. The behavioural recording is based on two distinctly colored, head-mounted LEDs, and the vector connecting the two LEDs determines the animal’s location (midpoint) and head direction. The behavioural data is recorded at 60 Hz, and downsampled to 30 Hz for all analysis.

In both tasks, no clustering is performed as we directly analyse sorted spikes used in the original study. Putative place cells and inhibitory interneurons are classified based on mean firing rates and spatial information of firing ratemaps (*58*) (Figure S3b). The LFP signal is retrieved from one representative electrode from one selected tetrode in both tasks.

##### Processing behavioural data

All second-order behavioural information, including speed, angular speed, and movement direction, are estimated from tracking data after temporal smoothing with a ±150 ms boxcar window. The allocentric goal direction is computed as the moment-by-moment angle prescribed by head location (midpoint of the vector prescribed by the LEDs) and the goal location. All relative directions of the form “*x* vs *y*”, are computed by setting the *y*-direction as the anchoring direction (0 degree), then computing the signed angle of the *x*-direction. For instance, “SD vs GD” indicates the relative sweep direction with respect to the goal direction (with goal direction being the 0 degree).

##### Processing spiking data

In the open-field experiment, hippocampal place cell data is constrained to periods above certain speed threshold (> 5 cm/s), to ensure robust theta rhythmicity and reduce noise in spatial tuning. By contrast, in the honeycomb maze task, the animal cannot move freely around the maze despite active engagement with the task contingency. We hence perform data exclusion when the following criteria are met simultaneously for a minimum duration of 50 ms: (a) theta power (6- 12 Hz) lower than 1 s.d. below the mean; (b) multi-unit activity greater than 2 s.d. above the mean; (c) ripple power (100-250 Hz) lower than 1 s.d. below the mean. The spectral power was computed as the magnitude of the Hilbert-transformed LFP signal.

#### Neural data analysis

##### Spatial decoding

Spatial decoding procedures for both tasks are largely similar, with only differences lying in the binning of the spatial enclosure. The two-dimensional spatial arena is partitioned into 40 × 32 bins (∼5 cm bin-size; *c.f*. 50 × 50 bins and 4 cm bin-size in the open-field experiment). The spatial tuning maps are computed as the smoothed (Gaussian-kernel, s.d., 8 cm) firing rates (number of spikes divided by the dwell time) within each spatial bin. To reduce falsely high estimates of firing rates, we post-process the ratemaps with Skaggs adaptive smoothing (*59*). Specifically, we place a circle centred at each bin location, and gradually increase the radius (in bins) such that the spikes/dwell-time within the expanded circle exceeds 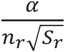, where *α* is a constant, *n*_*r*_ and *S*_*r*_ are the number of 50 ms-long occupancy samples and number of spikes lying within the circle with radius r, respectively. We set *α* = 2000 for both datasets. Only cells with peak firing rate (given the spatial tuning curves) > 1 Hz are selected in downstream spatial decoding. Width of place fields is computed as the radius of the largest circle around the peak firing location with all rates within the circle greater than 50% of the peak firing rate.

We estimate the rat’s spatial location based on a memoryless maximum-likelihood decoding algorithm (*22, 23*) (also known as the Bayesian decoder, assuming uniform prior), using the unit-specific spatial ratemaps and the population spike trains given temporal binning (which varies depending on specific analysis). Specifically, given a time window, Δ*t*, we assume each cell exhibits Poisson variability and the associated firing is exclusively modulated by spatial location. Leveraging Bayes rule, decoded spatial location is taken as the *maximum-a-posteriori* (MAP) estimate given the posterior distribution over spatial location.

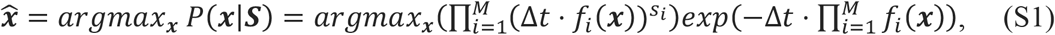

where *f*_*i*_(***x***) and *s*_*i*_ are the spatial tuning curve and number of emitted spikes within the selected time window of the i-th unit at location ***x***, respectively. Note that we have assumed independent firing rates across all neurons. Bayesian decoding yields accurate inference of the ground-truth spatial location, with high decoding accuracy on held-out validation sets given temporal binning of 250 ms (Figure S4). We only include putative place cells for spatial decoding, as the inclusion of inhibitory neurons lead to insignificant improvement in decoding accuracy (Figure S4b, c, f), largely due to the minimal gain in spatial information (Figure S3b). In addition to the MAP estimate, the full posteriors also reflect non-trivial uncertainty associated with spatial decoding. We hence additionally use the alternative estimate of the posterior mean to demonstrate the robustness of our findings (Figure S8).

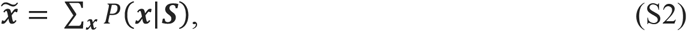

##### Theta sweep analysis

Theta cycles are retrieved from theta-filtered oscillations (6 – 12 Hz) of the local-field potential (LFP). The start and end of theta cycles are defined as the troughs of the Hilbert-transformed LFP. Leveraging the Bayesian decoder, decoded position within each theta sequence is estimated over a phase window of 60 degrees (equivalent to 14 - 28 ms), advancing in sliding phase window of 15 degrees (equivalent to 3.5 - 7 ms).

We additionally performed analyses for theta sweeps given theta cycles identified from the multi-unit activity (MUA; sampled at 30 kHz) given all recorded neurons (inclusive of interneurons), which is a smoothed histogram (Gaussian kernel, standard deviation of 5 ms) over time (sampled at 1 kHz). (Figure S9), using the similar approach based on Hilbert transform.

Notably, the sign of the recorded LFP activity can be arbitrary, and is dependent on the actual location of the tetrode (*60*). We hence apply constant 180-degree phase shift to all detected theta cycles given MUA such that the corresponding start and end times are maximally consistent with that of the LFP.

We observed that theta sweeps exhibit cyclic nature (Figure 1j), usually extending outward over one half of the cycle and progresses inwards to locations close to the animal over the other half. We hence only include the longest outward-extending subsequence (*9, 24*), lasting at least 180 degrees (half cycle), of each theta sweep in our analysis. Specifically, for each sequence, we fix the rat-specific start phase within the cycle (given the phase binning; honeycomb maze: 30 degrees for all rats in both honeycomb and open-field tasks), and select the end phase such that the resultant sequence length is longest. The initial and final offsets are taken to be the MAP estimates for the start and end phases of each sequence, and the corresponding sweep directions and lengths are computed as the allocentric angle and magnitude of vectors connecting the endpoints, respectively. To ensure robust theta modulation, theta cycles during which the rat is moving > 10 cm/s are included in the theta sweep analysis in the open-field task, whereas the speed threshold is set to 2 cm/s in the Honeycomb task due to the behavioural nature of the animal in the task configuration.

In plotting the roulette plot, we aggregated theta sweeps over equally sized angular bins (30 degree) of relative movement direction to the goal. Aggregation over sequence under varying rat poses in the environment requires standardising the decoded MAP estimates according to the moment-by-moment position and movement direction of the rat, through translation and rotations, which yields rat’s location at the origin, and movement direction pointing upwards post-standardisation.

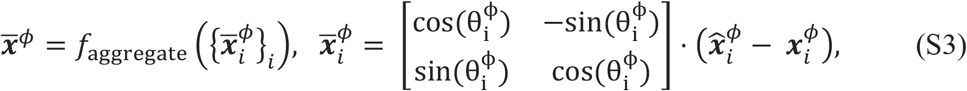

where 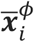 is the standardised MAP-estimate (post-translation and rotation), and 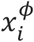 and 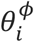 are the rat’s location and relative movement direction to the goal, at phase bin *ϕ* of the i-th sequence. In order to mitigate potential bias introduced by extreme events, the aggregation function, *f*_aggregate_(·), is taken to be the median decoded values along the orthogonal dimensions in the main paper. Note that aggregation with averaging lead to no significant differences in the resulting roulette plot (for both honeycomb and open-field tasks). Due to the varying lengths of sequences, sample size of the averaging varies within different phase bins. We found this leads to little effect in practice, due to the small variations in the lengths of selected theta sweeps (202.42 ± 17.02 degrees). We have also tried fixing the end phase for all sequences within each rat, which lead to identical findings qualitatively.

In the alternative analysis using posterior mean rather than MAP estimates for decoding theta sweeps (Figure S8c), we adopt a similar but subtly different aggregation procedure, through averaging over sequences before taking expectations. Specifically, given our spatial binning (40 × 32), we firstly embed the original posterior as the central block in an extended matrix of shape (120, 96) (three times the length of each side, to avoid losing information in subsequent translation and rotation transformations). We then translate the embedded posterior such that the rat’s location is placed at the centre of the matrix post-translation, followed by matrix rotation such that the movement direction is facing upwards (scipy.ndimage.rotate **(*61*)**). The aggregated estimate is then obtained by taking the expectation with respect to the averaging standardised posterior over all corresponding theta sweeps (Equation S2). The expectation is taken with respect to the central (40, 32)-block (normalised if necessary), since few decoded map locations lie greater than 1 m away from the animal’s location.

Quantitative comparisons of alignment between sweep directions and offset directions, and goal-, movement-, and head-directions are based on the Rayleigh concentration (or equivalently, the mean resultant vector length; MRL) of the circular distributions around its mean (which is usually close to 0 degree).

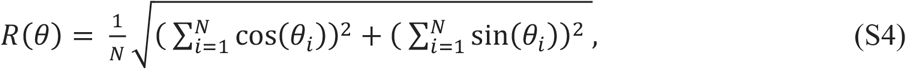

Where 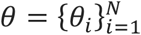 is the set of observations of the angular variable of interest.

We quantify the faithfulness of decoded theta sweeps with the quadrant score given the full decoding posterior (*37*). Specifically, given the projected theta reconstruction matrix over each theta cycle (Figure S9g, h), we focus on the central block that is within 100 cm of the animal’s location (at the beginning of the cycle), and within ± 90° of the cycle trough, and partition the enclosed area into 4 quadrants. Quadrants 2 and 4 represent probabilities associated with forward progression (given the projection), and quadrants 1 and 3 represent probabilities associated with backward progression. The quadrant score is then defined as the ratios between the sum of probabilities in quadrants 2 and 4 to the sum of probabilities in quadrants 1 and 3.

##### Replay sequence analysis

We restrict our detection of replay events within sharp wave/ripples, given the MUA of all clustered units. Events during which the speed is < 2 cm/s and the MUA greater than 3 s.d. from the mean, that last between 65 ms and 1000 ms are considered a valid candidate event for replay sequence decoding. Candidate events separated by less than 50 ms apart are merged and considered to be a single event. Candidate events during which the animal’s location is less than 50 cm from the goal location are excluded from analysis. Bayesian decoding of spatial locations (Equation S1) within replay events are based on temporal binning of 20-ms with 5-ms sliding window. Following (*21*), the replay sequence is truncated to the longest subsequence of time bins (but constrained to be > 50 ms) with peak posterior probability less than 20 cm from the preceding bin. We leverage similar standardisation procedures described above for aggregating theta sequences (Equation S3), but with additional scale-normalisation, such that the goal location is always placed 5 bins away from the animal’s location in the upward direction post- standardisation (similarly for the aggregation over the full decoding posteriors). For minimising random effects, we estimate the direction of replay sequences (RD) as the angle prescribed by the vector connecting the expected locations with respect to the averaged decoding posteriors aggregated over the first and second half of each candidate replay event. Replay sequence lengths are given by the lengths of the same vectors.

##### Phase coding analysis

To demonstrate reversed phase coding with respect to distance travelled along the trajectories, we constrained our analysis to periods of continuous movement towards or away from the goal, lasting greater than 800 ms (∼8 theta cycles) for robust examination of phase coding. Additionally, we allow sporadically brief discontinuity in the movement direction, by including trajectories with intermittent gaps < 150 ms of reversed direction, in order to maximise data utilisation. Normalised distance to the goal is computed as the path length along the trajectory (reversed when moving away from the goal), normalised between 0 and 1.

##### Statistical Analysis

Most statistical comparisons are on the rat-level, which we report the p-values of paired independent t-tests for assessment of statistical significance. We use two-sided tests unless there is strong prior belief about the relative relationship between the two groups under comparison (e.g., SD vs GD against SD vs MD). Some comparisons violate assumptions under t-tests, e.g., normality assumption for speed/angular-speed comparison due to the right-skewness, and unequal variances for comparison between theta sequences and replay sequences. In these cases, we use the Mann-Whitney U test, and report the statistical significance, as well as the effective size, r, as the significance can be deceiving especially given large sample sizes. Following the general wisdom, r < 0.2 is considered minor effects.

We use shuffled data to construct the null hypothesis distributions whereever necessary. In assessing the effect of goal-modulation in sweep direction in the honeycomb maze, movement modulation is a potential confound (despite absence of frequent goalward movement). Regressing out the effect of movement direction with circular regression (*25*), we then compute the MRL of the relative angles between residual sweep directions and goal-directions. We then shuffle the relationship between movement- and goal-directions, and perform the same residual analysis for each random shuffle, and compare the ground-truth MRL with the shuffled values.

#### Mechanistic modelling with continuous attractor networks

The computational model is a hierarchical system of continuous attractor networks, modelling theta-modulated para-subicular DCs as a ring-attractor, and entorhinal GCs and hippocampal PCs as two-dimensional attractors on a neuronal sheet.

Neurons in the DC attractor are organised over a ring, equally spaced between 0 and 360 degrees, with each cell active when the rat faces corresponding directions in the allocentric reference frame. The top-down goal-oriented directional input overrides the original internal directions driven by theta modulation and firing rate adaptation, yielding goalward internal direction sweeps.

Goalward directional sweeps drive activation of neurons in its downstream network of conjunctive GCs with preferred direction of firing along the directional sweep. The activity bump over the two-dimensional neuronal sheet in the conjunctive GC-network provide external phase-shift input to the GC-network, driving activity bump sweeping along the goal direction. The activity bump of the PC-network is then a feedforward projection of the GC activity bump. Due to the absence of periodic boundary condition in the PC-network (which is achieved effectively by instantiating significantly greater neuronal sheet), single-cell responses in the PC- network do not exhibit hexagonal firing fields, but retains goalward sweep arranged by theta cycles.

Below, we provide mathematical details of each sub-component of the model. Parameters used in all simulations (Figure 3, Figure S13) can be found in Table S1. All simulations are numerically integrated with exponential Euler method (*62*), and were implemented using the BrainPy library (*63*).

##### Directional ring attractor

Directional cells (e.g., subicular head-direction cells) are assumed to lie on a ring attractor, receiving top-down goal-oriented directional inputs, and is modulated by theta rhythmicity and exhibits firing rate adaption mechanism (*9, 64*).

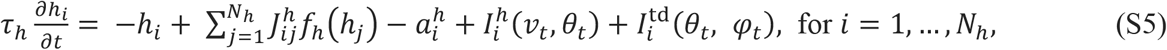

where *τ*_*h*_ is the time constant, *h*_*i*_ is the pre-synaptic input to the i-th directional cell and *N*_*h*_ is the number of neurons in the DC network, 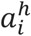 denotes the corresponding firing rate adaptation modulation, 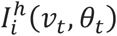 denotes the directional-dependent external sensory inputs that is jointly modulated by speed and theta rhythmicity, 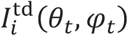, denotes the top-down goal-modulated directional input, and *v*_*t*_ *θ*_*t*_ and *φ*_*t*_are the animal’s speed, allocentric head direction, and goal direction, respectively.

The external sensory input is modelled with a squared-exponential response function, with peak at animal’s current head direction, *θ*_*t*_.

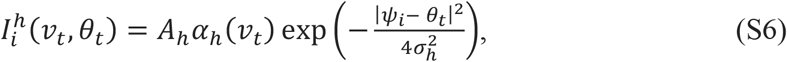

where *ψ*_*i*_ represents the preferred internal direction of i-th neuron (in radians), |*ψ*_*i*_ − *θ*_*t*_|^2^ represents circular distance between the internal direction for the i-th neuron and current head direction, *σ*_*h*_ controls the tuning width of sensory input, *A*_*h*_ denotes the strength of sensory input, *α*_*h*_(*v*_*t*_) denotes the speed-regulated theta modulations from medial septum (*1*), in the form of sinusoidal waves.

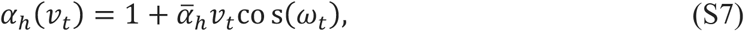

where 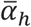 denotes the strength of theta modulation, and *ω*_*t*_ is the theta phase at time t. Note that we assume the preferred internal directions of neurons in the DC network tile the [0, 2π] space with equal spacings.

The top-down goal-oriented directional input, potentially originated in the RSC, is also modelled with a squared-exponential tuning, with peak at the allocentric goal direction, *φ*_*t*_.

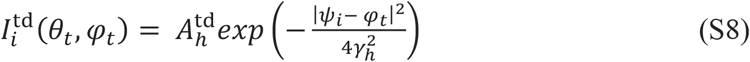

where 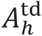 and 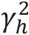 denotes the strength and tuning width of top-down directional inputs, respectively.

The non-linear activation function is modelled as second-order global inhibition.

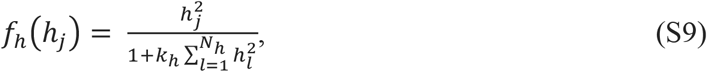

where *k*_*h*_ represents the global inhibition strength within the DC network. Previously works have shown global inhibition is crucial for maintaining localised activity bump on the ring attractor (*12, 65*).

The recurrent connections between neurons i and j, with preferred firing directions *ψ*_*i*_ and *ψ*_*j*_ (in radians), respectively, is defined by a circular squared exponential connectivity.

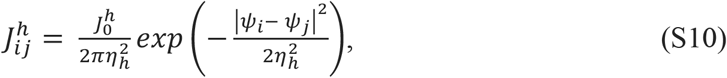

where 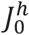 and *η*_*h*_ control the strength and tuning widths of the DC connectivity, respectively.

The firing rate adaptation, 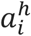, is implemented to mechanistically model the negative feedbacks that induces dynamic reduction in neuron’s firing rate in response to constant-intensity inputs (*13, 18*).

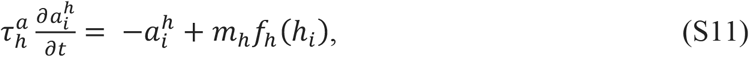

where 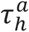 is the time constant of firing rate adaptation, *m*_*h*_ represents the adaptation strength. The time constant, 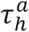, is usually set to be a magnitude greater than *τ*_*h*_, modelling the effect that firing rate adaptation is a slower dynamics compared to spikings. At the network level, firing rate adaptation induces non-trivial intrinsic dynamics to the activity bump, leading to sequential activation of neurons in the absence of sensory updates.

###### Place cell attractor network

Place cells are assumed to lie on a two-dimensional neural sheet, with equally spaced field centres. The PC network receives spatial sensory inputs, and theta-modulated phase-offset inputs from conjunctive GCs, with the associated activity driven by the projection of DC activities.

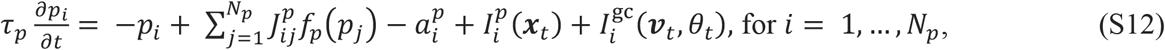

where *τ*_*p*_ is the time constant, *p*_*i*_ is the pre-synaptic input to the i-th place cell, *N*_*p*_ is the number of place cells, 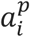 denotes the firing rate adaptation, 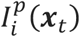 denotes the spatially selective external sensory input at location ***x***_*t*_ and 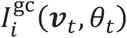 denotes the directionally-shifted input from conjunctive grid cells with joint modulations from theta oscillations and speed.

We assume global inhibition non-linear activities in place cells are of the same form as head direction cells (Equation S9), but with different inhibition strength, *k*_*p*_, and additional population-wide gain factor, *g*.

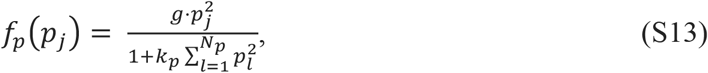

Similarly, we assume Gaussian-like recurrent connections between place cells I and j, with field centres ***s***_*i*_ and ***s***_*j*_.

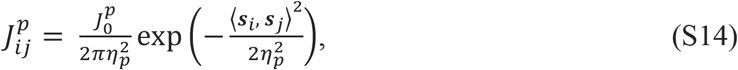

where ⟨***s***_*i*_, ***s***_*j*_⟩ denotes the Euclidean distance between field centres ***s***_*i*_ and ***s***_*j*_, 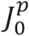 and *η*_*p*_ control the strength and tuning widths of the place cell connectivity, respectively.

The strength of external sensory input is also modelled with Gaussian tuning, with peak at the allocentric spatial location, ***x***_*t*_.

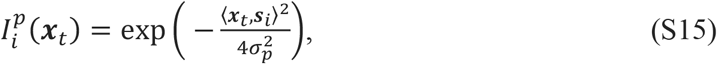

We omit explicit modelling of grid cells in the current model, due to GPU-memory constraints (given the modelling of conjunctive grid cells in all directions, and canonical grid cells in different modules). Instead, we model the effect given projections from upstream conjunctive grid cells in the simplified form of directionally shifted spatial inputs to the place cells. Assuming surjective Hebbian plasticity from entorhinal grid cells to CA1 place cells with overlapping firing fields, the direct consequence of shifted spatial inputs, such that grid-cell activity bumps shift along the goal direction, is effectively translated to goalward sweeping of place cell activity bumps, hence yielding goal-directed hippocampal theta sweeps.

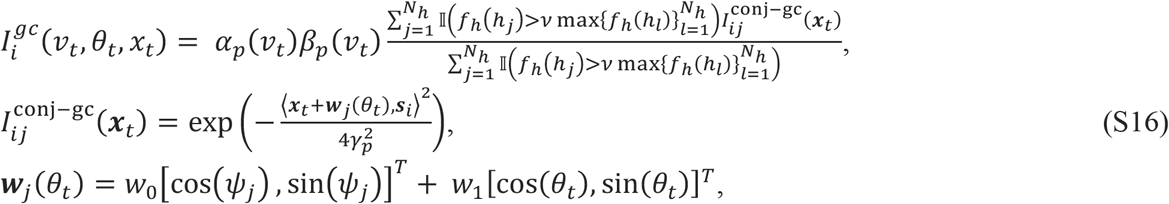

where the strength of input scales linearly with running speed.

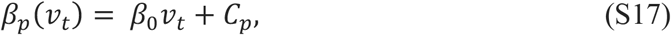

for constants *β*_0_ and 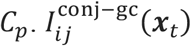 represents the Gaussian-shaped projection strength from the phase-shifted activity bump of conjunctive grid cells with preferred internal direction *ψ*_*j*_, at the current location ***x***_*t*_. The phase shift is towards both the internal direction and current head/movement direction. 𝕀(·) is the indicator function, modelling a soft winner-takes-all thresholding mechanism, given ν ∈ [0, 1] that sets the threshold on firing rates of head direction cells for selecting the subset of idealised conjunctive grid cells to activate, ensures only conjunctive grid cells with preferred internal direction lying along the current goal direction (modulated by the head direction cell activity) contributing to the construction of directionally- shifted spatial input. This effectively shifts the place cell activity bump in the direction of the goal. We additionally induce speed-regulated septal theta-modulation, *α*_*p*_(*v*_*t*_), given that conjunctive grid cells are tuned to theta rhythm (*27*).

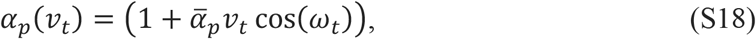

where 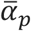 is the strength of theta modulation.

Place cells follow the same firing rate adaptation mechanism as the grid cells, with time constant, 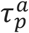, and adaptation strength, *m*_*p*_.

##### Simulated trajectory

Simulated trajectories aim to capture the dominant features in rat’s real behaviour in the honeycomb maze (Figure S6a). For simplicity, we model the rat body as a rigid stick, with the outward end representing the head location, which enables efficient simulation of rotational scanning movements. We interleave random walk and random head-rotation movements, with constant speeds. At each timepoint (simulated at 1000 Hz), with 0.5 probability, the simulated rat engages in rotational scanning movement spanning angles sampled uniformly from [0.3π, π] (Figure 3c).

##### Theta sweep decoding and phase coding with mechanistic simulations

The proposed CAN is rate-based by design, and instead of instantiating a spiking model for decoding theta sweep based on generated spikes, we use the peak location of the place cell activity bump as a proxy for the decoded location, which is unbiased on expectation.

For simulating phase precession/procession, we simulate Poisson spikes given the instantaneous firing rates of each place cells. To reduce cluttering in visualisation, we apply a gain factor of 0.05 to place cells firing rates to reduce the total number of spikes emitted.

**Figure S1.**
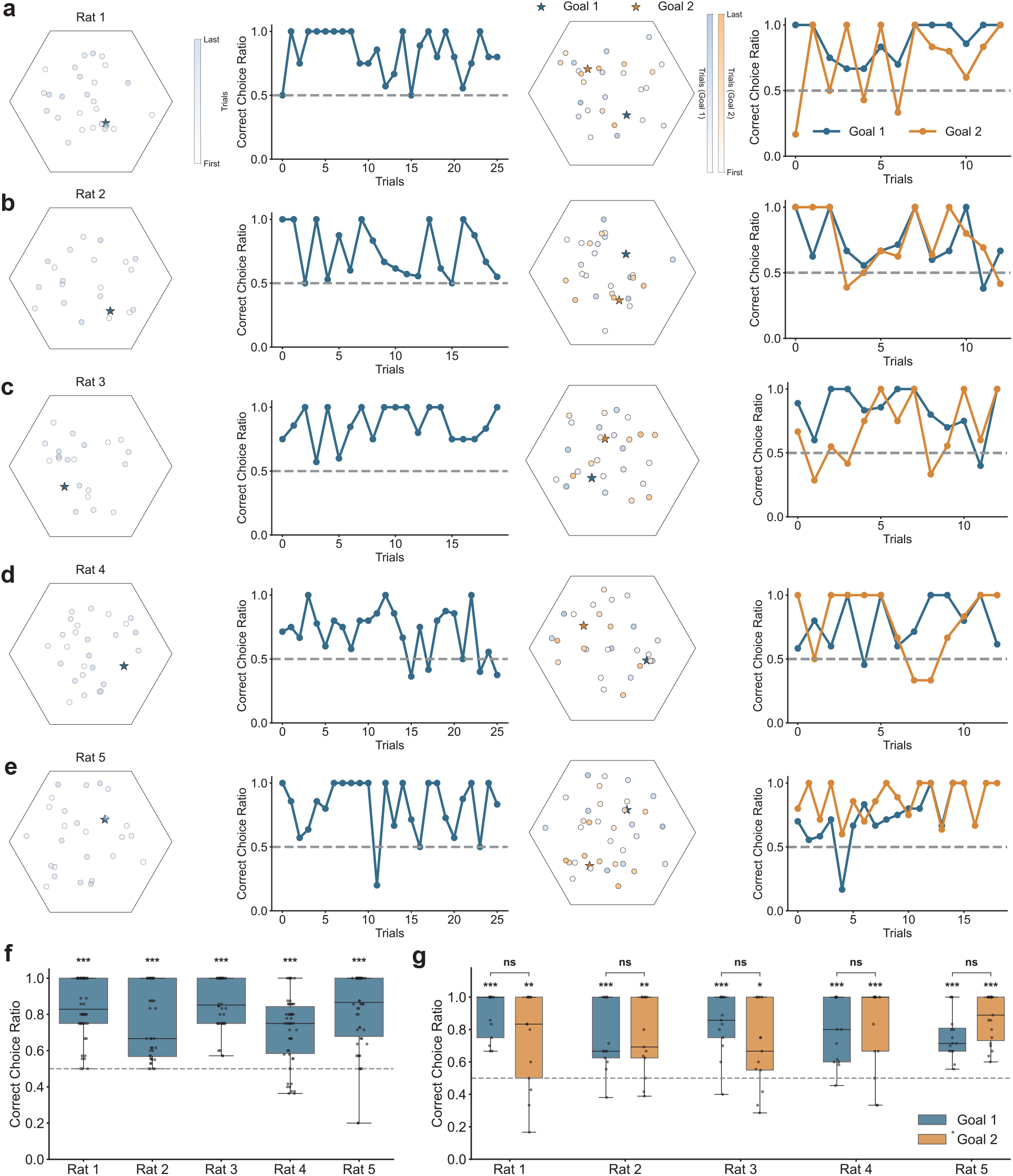
Rats learned to correctly navigate to the goal in the Honeycomb maze. **a**. For Rat 1, from left to right: goal location and trial-specific start locations in the single-goal session; trial-specific ratio of correct choices; goal locations and corresponding trial-specific start locations for both goals in the goal-switching session; trial-specific ratio of correct choices before and after the goal switching. Note that trials are in block order and the goal-switching only happens once. **b-e**. Same as (a), but for Rats 2-5. Note that Rat 5 was trained over extended number of training sessions, and here we present the behavioural data from the final session. **f**. Proportions of trial-specific correct choices for each rat in the single-goal session (one-sided one-sample t-test between the trial-specific correct choice ratios and chance level; Rat 1: p = 1.6600 × 10^−10^; Rat 2: p = 1.3010 × 10^−5^; Rat 3: p = 3.5644 × 10^−10^; Rat 4: p = 3.1467 × 10^−6^; Rat 5: p = 3.8588 × 10^−8^). **g**. Same as (f), but separately for the two goal locations in the goal-switching session (two-sided paired t-test between the trial-specific correct choice ratios for the two set of trials corresponding to different goals; Rat 1: p = 0.1094; Rat 2: p = 0.5647; Rat 3: p = 0.1082; Rat 4: p = 0.5544; Rat 5: p = 0.0541). Note that Rat 5 is excluded from all analyses associated with goal-switching experiments.

**Figure S2.**
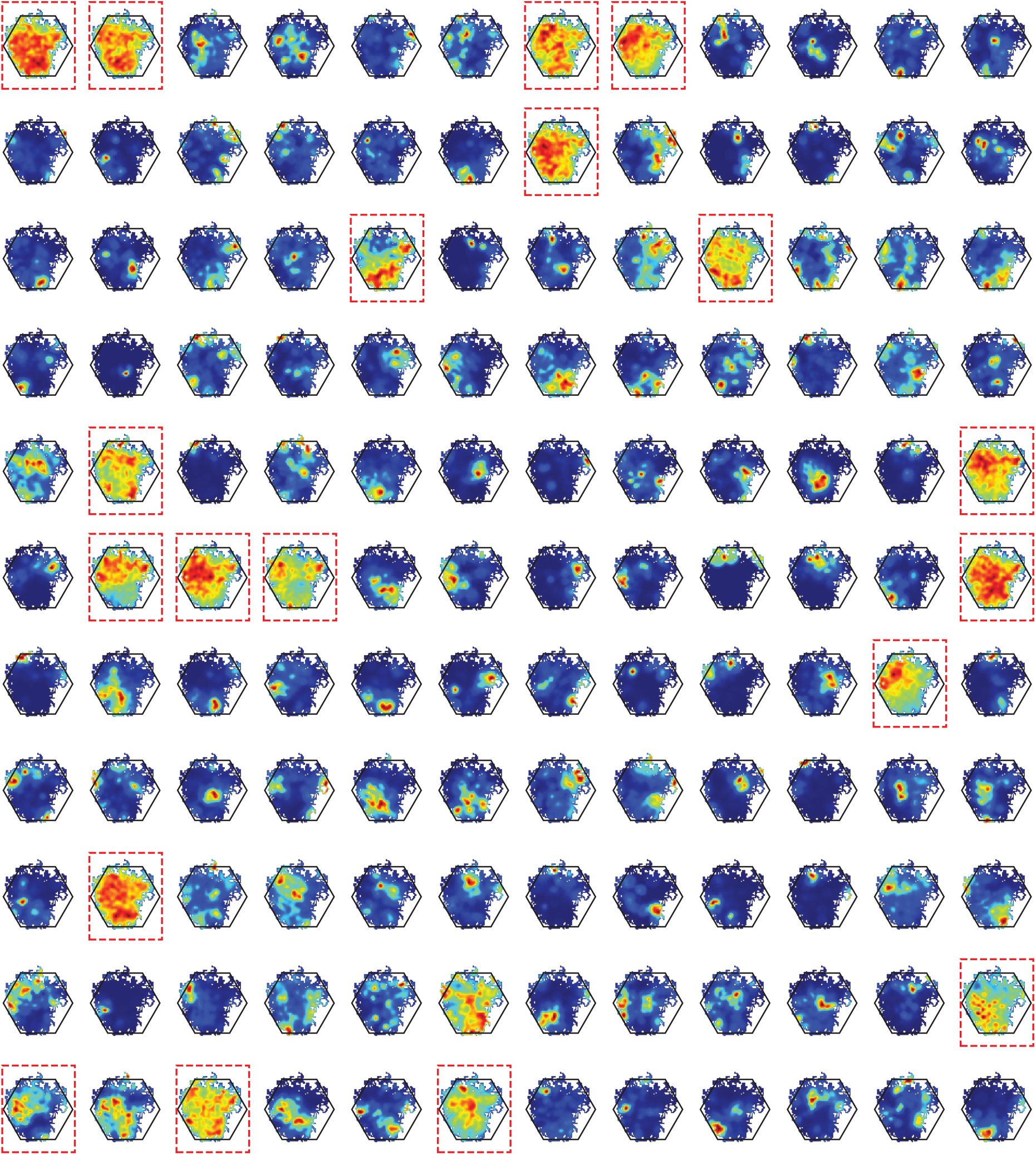
Spatial tuning for all recorded neurons in Rat 2, with Skaggs smoothing (*59*) Cells enclosed by the red rectangular box are classified as interneurons (19 / 132, see Methods), and rest are putative place cells.

**Figure S3.**
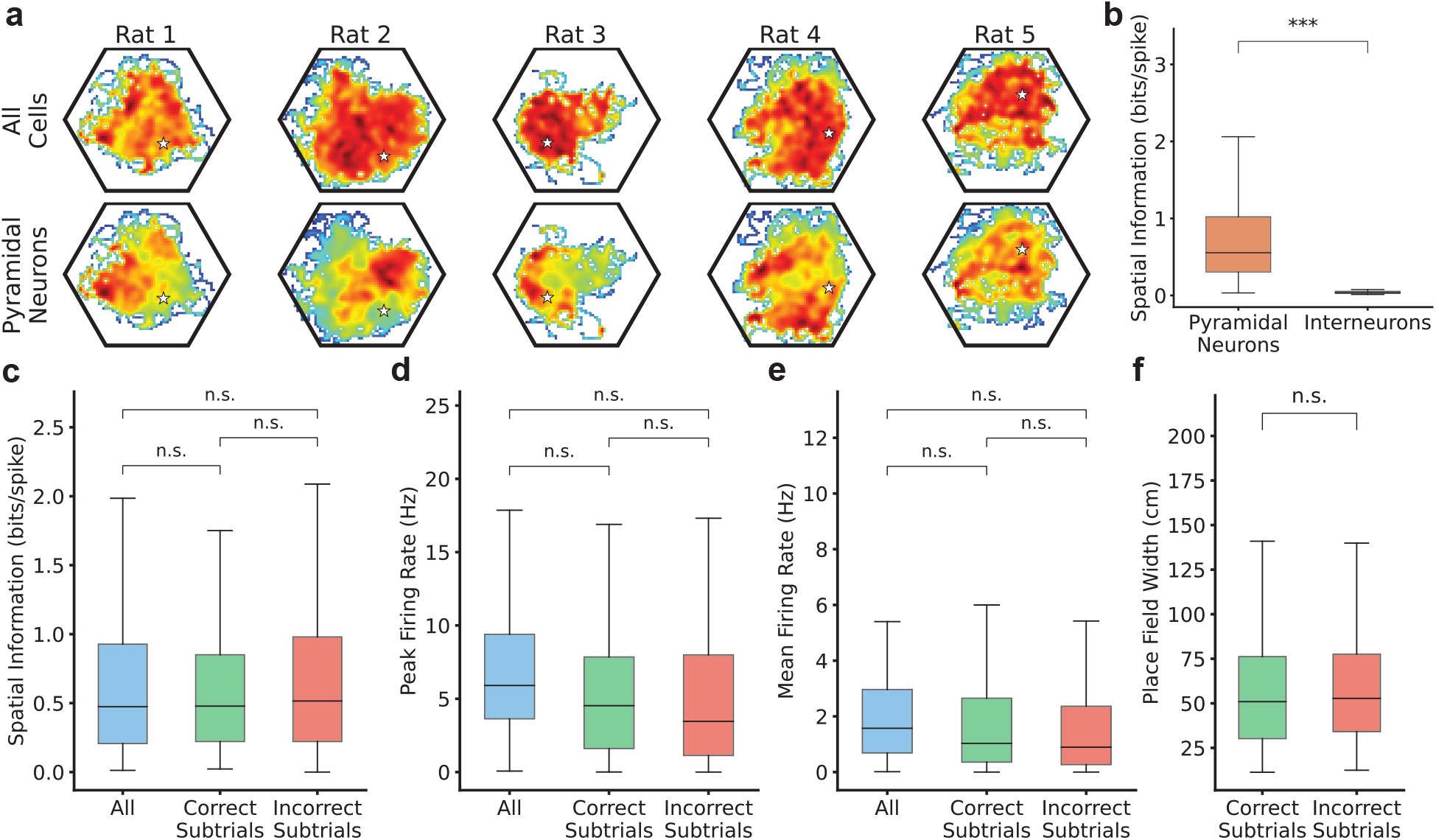
Properties of place cells firing in the Honeycomb maze. **a**. Rat-specific aggregated spatial tuning (summation of all cell-specific spatial tuning curves) for all recorded neurons (top), and all putative pyramidal neurons(bottom). White star indicates the goal location. **b**. Cell-specific spatial information (bits per spike (*58*); see Methods) for pyramidal neurons and interneurons (one-sided Mann-Whitney U-test: p < 0.001, effective size: r = 0.9558). There is little spatial information inherent in inhibitory interneurons, we hence expect the inclusion of interneurons for spatial decoding would lead to minimal improvement in terms decoding accuracy (apart from Rat 3, also see Figure S4} for empirical evidence). **c**. Cell-specific spatial information over all valid periods, and “wait-periods” preceding correct and incorrect choices, respectively, for all recorded neurons (paired t-test; all vs correct choices: p = 0.0780, all vs incorrect choices: p = 0.9363, correct vs incorrect choices: p = 0.8058). **d-e**. Same comparison as (c), but for cell-specific peak firing rates (d; two-sided paired t-test; all vs correct choices: p = 0.2583, all vs incorrect choices: p = 0.1427, correct vs incorrect choices: p = 0.0997), and mean firing rates over all binned spatial locations (e; two-sided paired t-test; all vs correct choices: p = 0.4793, all vs incorrect choices: p = 0.1123, correct vs incorrect choices: p = 0.2185). **f**. Place field width (see Methods) for all pyramidal neurons over correct and incorrect choices (two-sided paired t-test; p = 0.9047). We do not find any significant difference between firing properties of place cells under correct or incorrect choices, we hence include all valid periods in recorded trials, regardless of the choice correctness, in our spatial decoding and sweep analysis.

**Figure S4.**
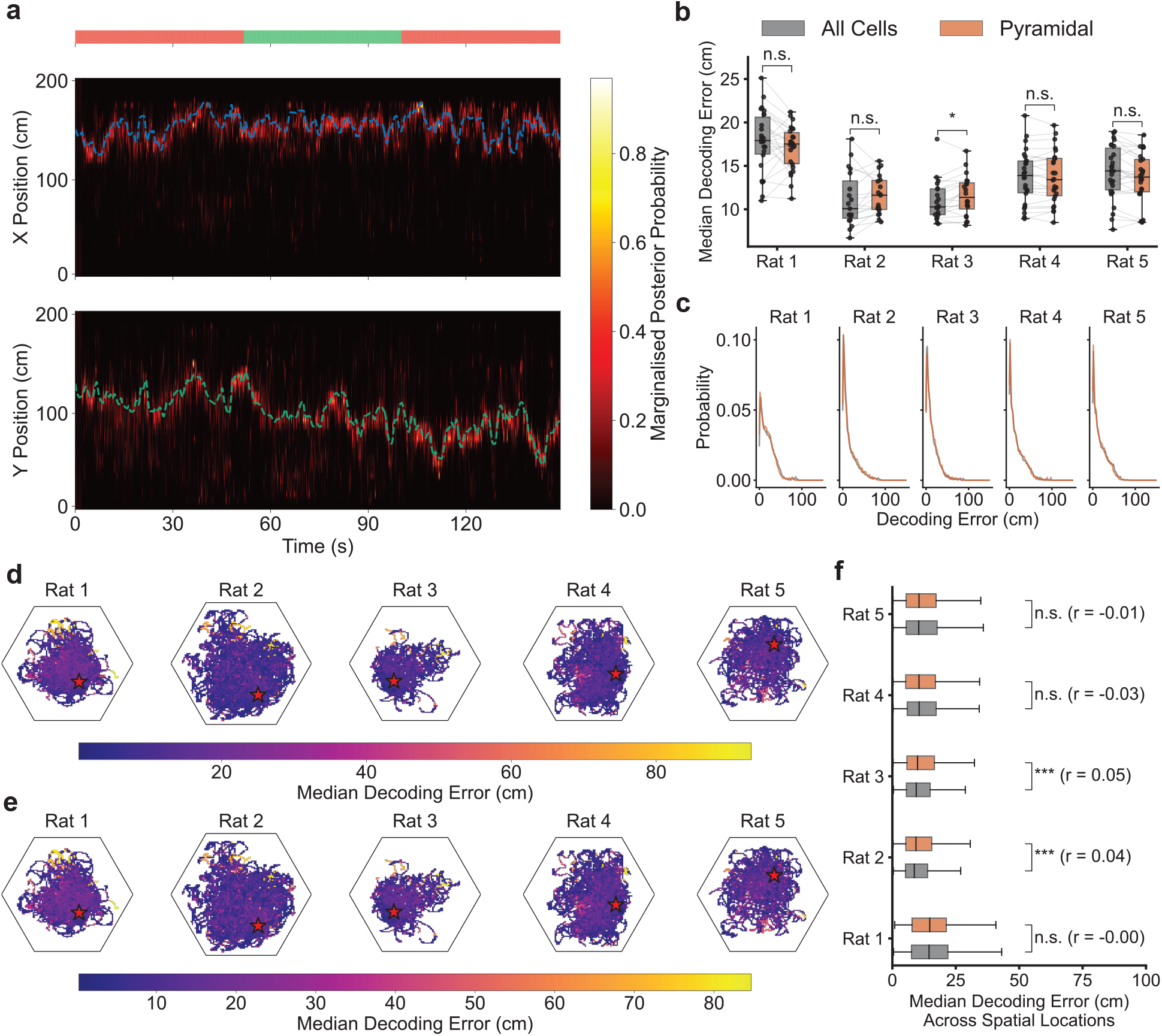
Bayesian decoding of spatial location in the Honeycomb maze. **a**. Exemplary posterior probabilities over a held- out validation trajectory spanning 145 seconds given Bayesian decoding, plotted separately for the × (top) and y (bottom) positions (Rat 2). The probabilities are computed via marginalisation over the orthogonal dimension. Dashed lines represent the ground-truth locations of the rat (blue for x-position and green for y-position). The horizontal bar at the top indicates the choice- correctness of the ongoing “subtrial” (green and red indicate correct and incorrect choices, respectively). Qualitative inspection shows that Bayesian decoding yields accurate inference of the rat’s spatial position (median decoding error: 13.1084±.5635 cm, 5 rats). **b**. Median decoding errors within each trial, using either all cells (grey) or only pyramidal cells (orange), for all 5 rats (two-sided paired t-test between the two groups of decoding errors; Rat 1: p = 0.9232, Rat 2: p = 0.1319, Rat 3: p = 0.0154, Rat 4: p = 0.6799, Rat 5: p = 0.9941). Note that decoding errors are generally low for both approaches, with the exception of rat 3, where decoding with all neurons leads to lower errors than the alternative approach. This is likely due to the overall insufficient coverage of the spatial environment given recorded pyramidal neurons (Figure S3), yielding more accurate spatial decoding given additional spatial information from interneurons. **c**. Distribution of all decoding errors (aggregating over all trials) given Bayesian decoders with either all cells (grey) or only pyramidal cells (orange). **d**. Median decoding error over spatial locations in the Honeycomb maze, for all 5 rats, given a Bayesian decoder with all cells. Goal location is indicated by the red star. **e**. Same as (d), but using a Bayesian decoder with only pyramidal neurons. **f**. Comparison of median decoding error over spatial locations between the two Bayesian decoders (One-sided Mann-Whitney U test; Rat 1: p = 0.5946, Rat 2: p = 0.0002, Rat 3: p = 0.0008, Rat 4: p = 0.9722, Rat 5: p = 0.6795), along with corresponding effective size of the test (r). Across trials and spatial locations, the difference in decoding accuracy between Bayesian decoders given all cells and only pyramidal cells is minor, we hence proceed with the latter in all analyses in the paper.

**Figure S5.**
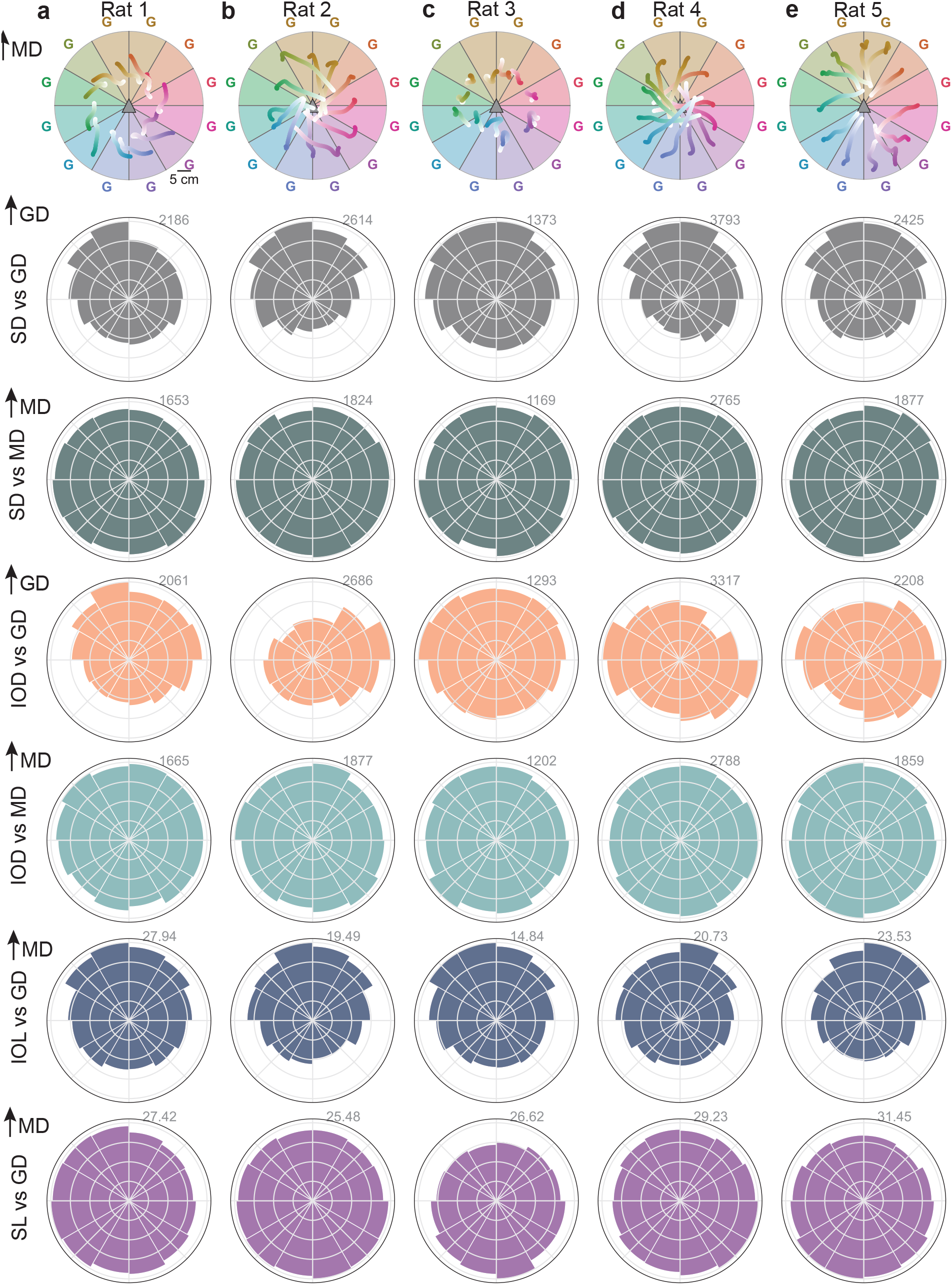
Goal-oriented theta sweeps in individual rats. **a**. Rat 1. From top to bottom: median theta sweeps as a function of relative movement direction (upward) to the goal; circular histogram of relative angles between sweep- and goal-directions; circular histogram of relative angles between sweep- and movement-directions; circular histogram of relative angles between initial offset- and goal-directions; circular histogram of relative angles between initial offset-direction and movement-direction; median lengths of initial offset of decoded theta sweeps given varying relative movement direction to the goal; median lengths of theta sweeps given varying relative movement direction to the goal. **b-e**. Same as (a), but for Rats 2 - 5. Data from individual rats yield similar qualitative observations as in the aggregated analysis (Figure 1): theta sweeps within individual rats exhibit strong goal-oriented direction bias.

**Figure S6.**
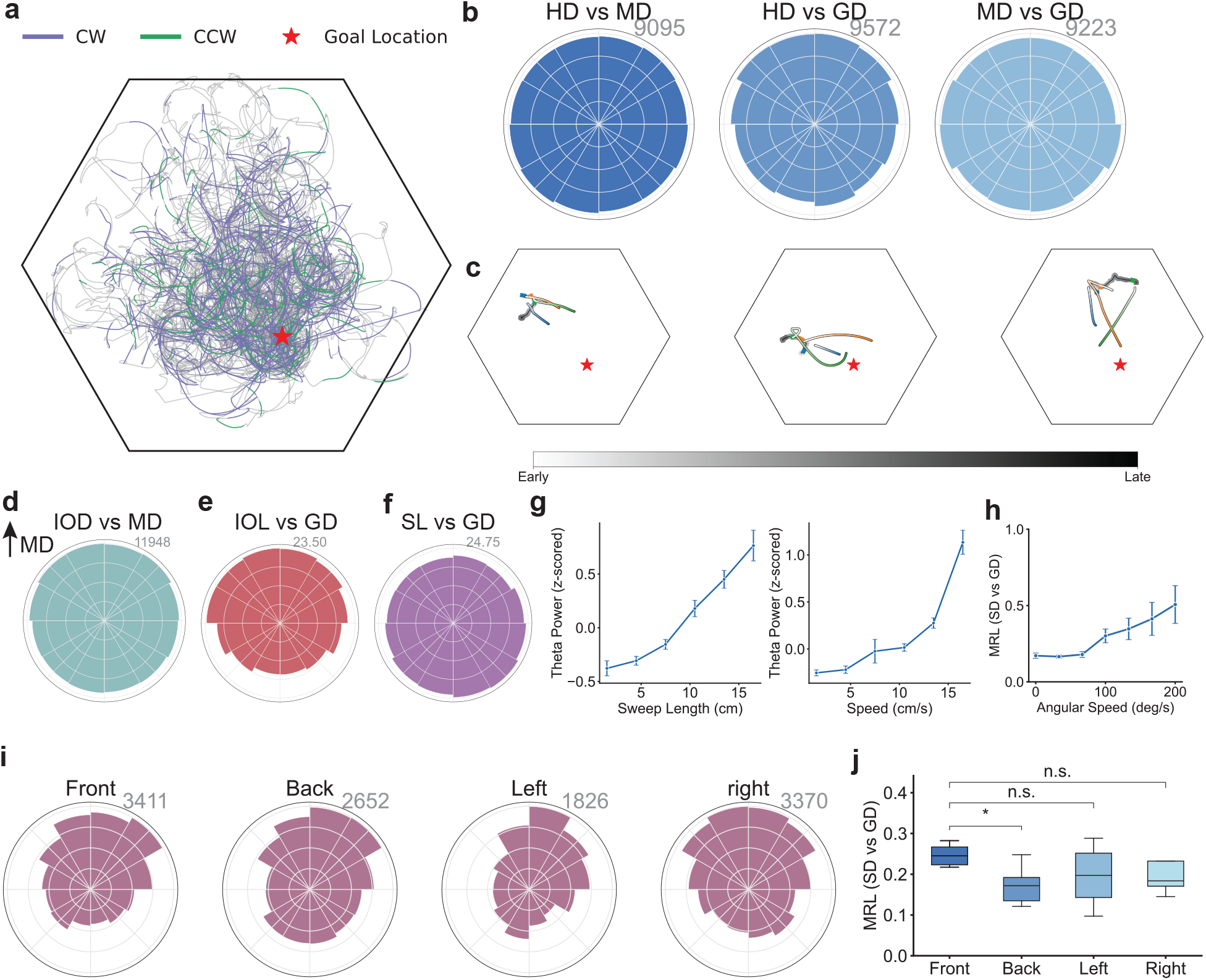
Theta sweeps exhibit strong goal-oriented and moderate movement-oriented directional bias in the honeycomb maze. **a**. Position trace over all periods (gray) and over actively scanning periods (magenta and green for CW- and CCW- rotations, respectively), over all trials within the single-goal session of Rat 1. **b**. Circular histograms of relative angles between HD, MD, and GD in the honeycomb maze, aggregated over all rats. The behavioural paradigm effectively reduces goalward movement and headings. **c**. Demonstrations of goal-oriented theta sweeps over three trajectories with different relative movement directions to the goal (gray line). Decoded theta sweeps and corresponding locations over concurrent movement are shown in matching colors. Goal location is indicated by the red star. **d**. Circular histogram of relative angles between IOD and MD. **e**. Median lengths of initial offset of decoded theta sweeps (relative to rat’s location) given varying relative MD to the goal (movement direction is upwards). **f**. Same as (e), but for sweep lengths. **g**. Theta power (z-transformed) as a function of increasing sweep lengths (left) and movement speed (right). **h**. MRL of the distribution of relative angles between SD and GD as a function of increasing angular speed (of head rotations). **i**. Circular histograms of relative angles between SD and GD (from left to right) when rats are moving towards, opposite to, to the left and right of the goal direction, respectively (aggregating over all rats; goal direction is upwards). **j**. Rat-specific MRL of the distributions in (i) (two-sided paired t-test, Front vs Back: p = 0.0106; Front vs Left: p = 0.1632; Front vs Right: p = 0.1005). Directional bias in theta sweeps is jointly modulated by the movement and goal direction (as indicated by the Front-Back comparison in i and j), and the goal modulation is significantly stronger (Figure 1g, i).

**Figure S7.**
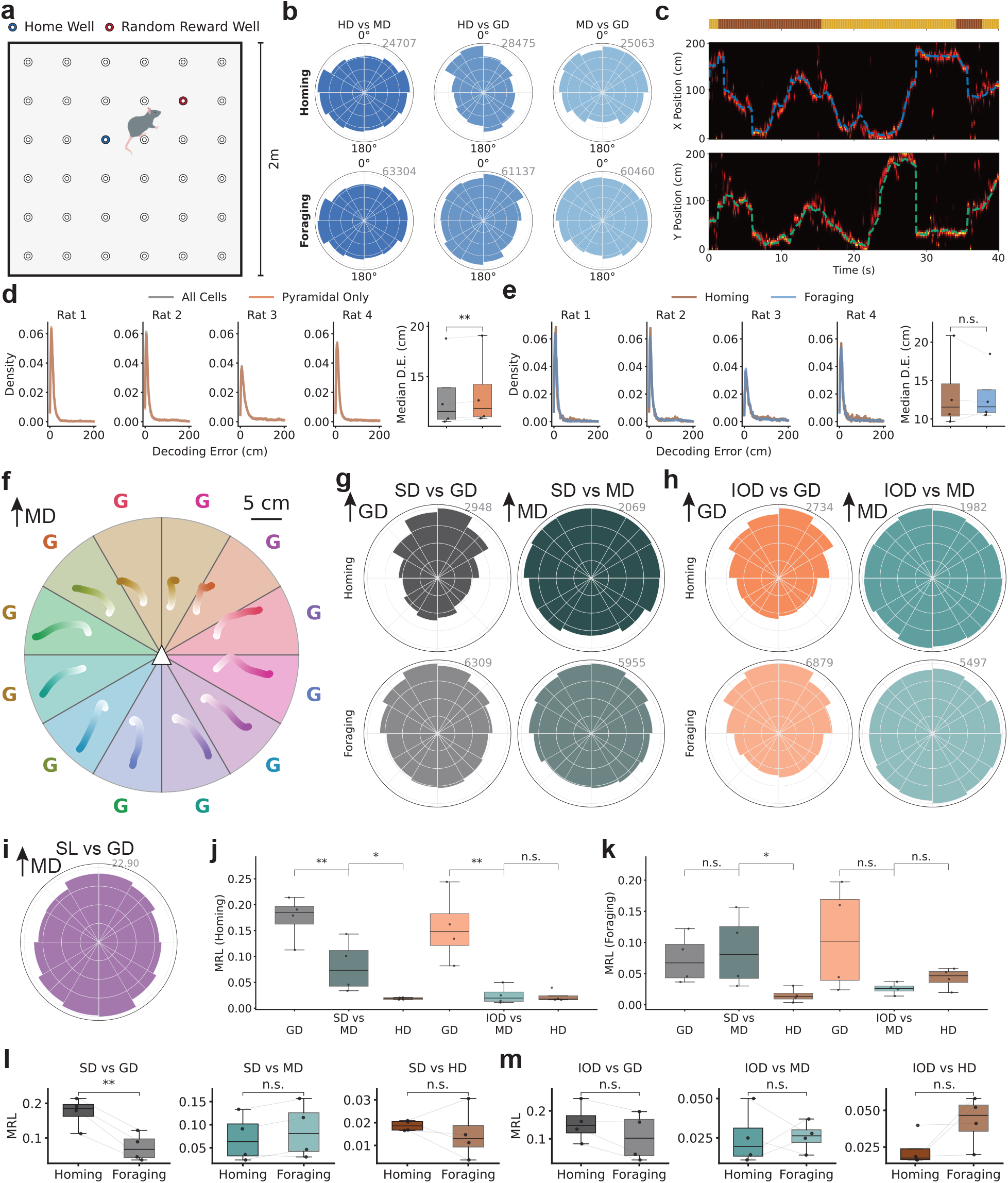
Theta sweeps exhibit goal-oriented directional bias in spatial memory tasks in open-field environments. **a**. Graphical demonstration of the open-field spatial memory task from (*21*). Within each session, rats alternate between “*homing*” trials, during which rats need to navigate to a fixed home well, and “*foraging*” trials, during which rats explore the environment to find rewards in a randomly selected remote well (see Methods). **b**. Circular histograms of relative angles between (from left to right) HD and MD, HD and GD, MD and GD, over homing (top) and foraging (bottom) trials. Rats engaging in spatial navigation tasks in open-field environments exhibit strong alignment between head/movement direction and GD (*c.f*. Figure S6b). Goal- movement alignment is stronger over homing trials compared to foraging trials. **c**. Example Bayesian-decoding posterior probabilities over a held-out validation period lasting 40 seconds, plotted separately for the x- (top) and y- (bottom) positions. The presented probabilities are marginalised over the orthogonal dimension. Dashed lines represent the ground-truth x- (blue) and y- (green) locations of the rat. The horizontal bar at the top indicates the trial identity (brown for homing trials and yellow for foraging trials) over the selected period. Qualitative inspection show that Bayesian decoding yields accurate inference of the rat’s spatial position (median decoding error: 13.1050 ± 0.3490 cm, 4 rats). **d**. Probability of decoding errors given all cells and only putative pyramidal neurons for Rats 1-4 (from left to right). Rightmost panel: rat-specific median decoding error given Bayesian decoding using all cells and only pyramidal neurons (two-sided paired t-test: p = 0.0012). Unlike spatial decoding in the honeycomb maze (Figure S4), using all cells (including inhibitory interneurons) yields spatial decoding significantly more accurate across all rats. We hence proceed with such choice in associated analysis. **e**. Probability density of decoding errors over homing and foraging trials for Rats 1-4 (from left to right). Rightmost panel: rat-specific median decoding error over homing and foraging trials (two-sided paired t-test: p = 0.6622). No significant difference in decoding accuracy is observed between homing and foraging trials. **f**. Median theta sweeps as a function of varying relative movement direction to the goal (indicated by coloured “G”s). **g**. Circular histograms of relative angles between SD and GD (left), and SD and MD (right), over homing (top) and foraging (bottom) trials. **h**. Same as (g), but for the direction of initial offsets of theta sweeps. **i**. Median sweep lengths as a function of varying relative movement direction (upward) to the goal. **j**. Comparison of rat-specific MRL of the empirical distributions in (g) and (h), during homing trials (one-sided paired t-test; wrt SD, GD vs MD: p = 0.0071; two-sided paired t-test; MD vs HD: p = 0.0451; wrt IOD, GD vs MD: p = 0.0079, MD vs HD: p = 0.2889). **k**. Same as (i), but over foraging trials (one-sided paired t-test; wrt SD, GD vs MD: p = 0.6445, two-sided paired t-test; MD vs HD: p = 0.0270; wrt IOD, GD vs MD: p = 0.0546, MD vs HD: p = 0.9341). Analysing theta sweeps in the open-field task yields same main findings as in the honeycomb maze analysis (*c.f*. Figure 1): theta sweep directions are jointly modulated by goal and movement directions; with goal modulation significantly stronger than movement modulation, directions of initial offsets are exclusively modulated by goal direction; theta sweeps extend longer when rats are moving away from the goal. **l**. Comparison of rat-specific MRL of the empirical distributions in (j) and (k), over homing and foraging trials (one-sided paired t-test; SD vs GD: p = 0.0018; two-sided paired t-test; SD vs MD: p = 0.1082; SD vs HD: p = 0.5234). **m**. Same as (l), but for the IOD-related empirical distributions in panel g (two-sided paired t-test; IOD vs GD: p = 0.0730; IOD vs MD: p = 0.9373; IOD vs HD: p = 0.1629). We observe stronger goal modulation and similar-level movement modulation in homing trials, whereas both modulations are of similar strength during foraging trials. The differential modulatory profiles between homing and foraging trials further substantiates the validity and cognitive affordance of goal-oriented directional bias in theta sweeps.

**Figure S8.**
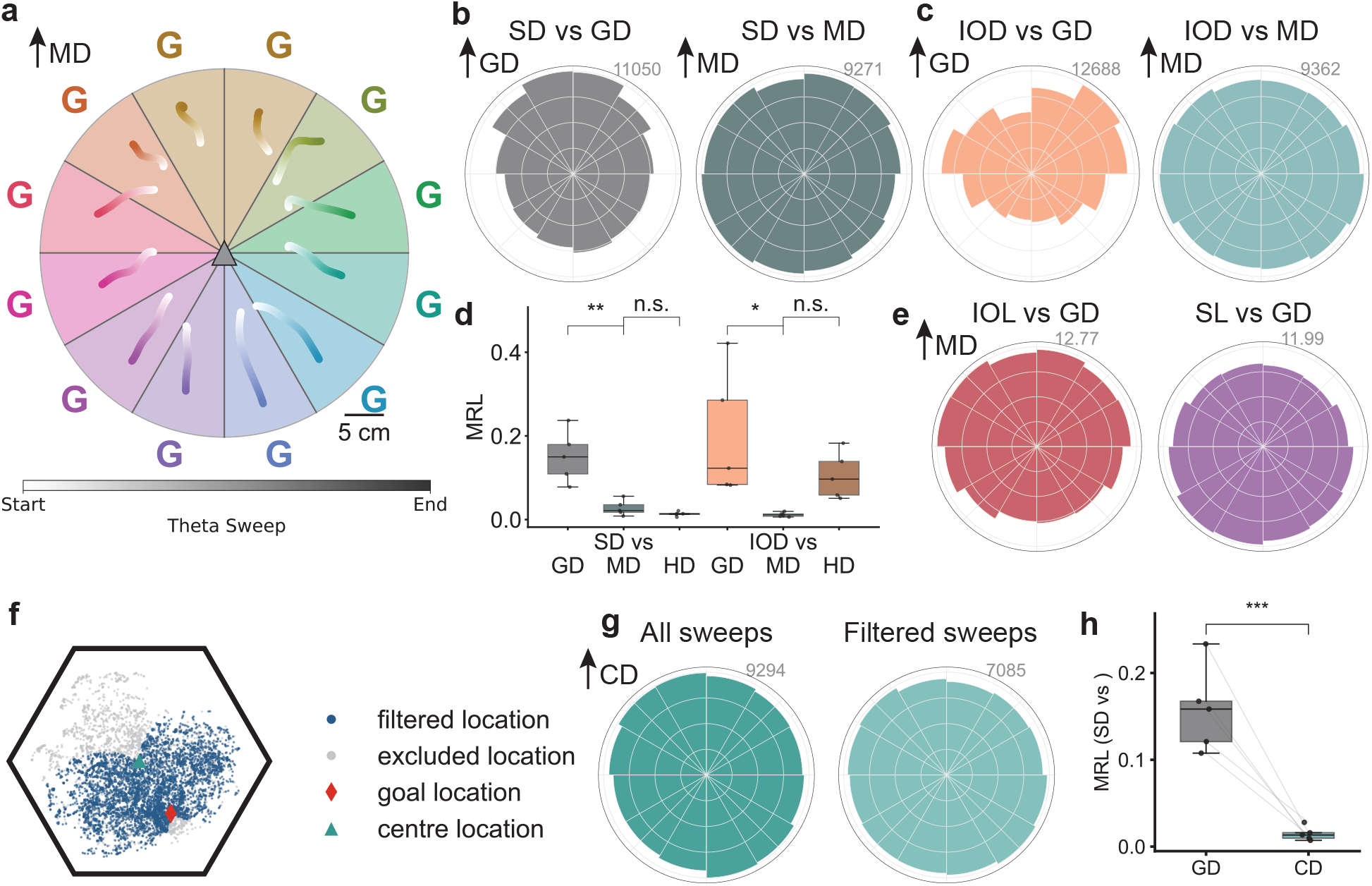
Goal-oriented directional bias in theta sweeps using decoding distributions. Theta sweeps are defined by the decoded locations, i.e. choosing the *maximum-a-posteriori* estimate from the probability distribution at each timestep, and these are aggregated across trials. Here we aggregated the full distributions rather than taking the single most likely location. **a**. Median theta sweeps as a function of varying relative MD to the goal (MD upwards). **b**. Circular histograms of relative angles between SD and GD (left), and SD and MD (right). **c**. Same as (b), but for the IOD. **d**. Comparison of rat-specific MRL of the circular distributions in (b) and (c) (one-sided paired t-test SD wrt GD vs MD: p = 0.0014; two-sided paired t-test SD wrt MD vs HD: p = 0.0671; IOD wrt GD vs MD: p = 0.0116; IOD wrt MD vs HD: p = 0.9973.). **e**. Median distance from initial offset of decoded theta sweeps to rat’s location (IOL; left), and median sweep lengths (SL; right), as a function of movement direction relative to goal direction (MD upwards). **f**. Demonstration of control for centroid bias. Taking expectations with respect to the posterior distribution will inevitably yield decoded locations biasing towards the centre of the environment. Here we control for such bias and remove theta sweeps with the corresponding rat’s location (approximately) lying along the extension of the line connecting the goal and centre locations (± 30°). Additional analysis on filtered sequences removes potential confound arising from overlapping directions to the goal and centre. **g**. Histograms of relative angles between sweep direction and centroid direction (CD), over all (left) and filtered (right) theta sweeps. **h**. Rat-specific MRL of relative sweep direction with respect to the goal direction and centroid direction, over filtered theta sweeps (one-sided paired t-test: p = 0.0001).

**Figure S9.**
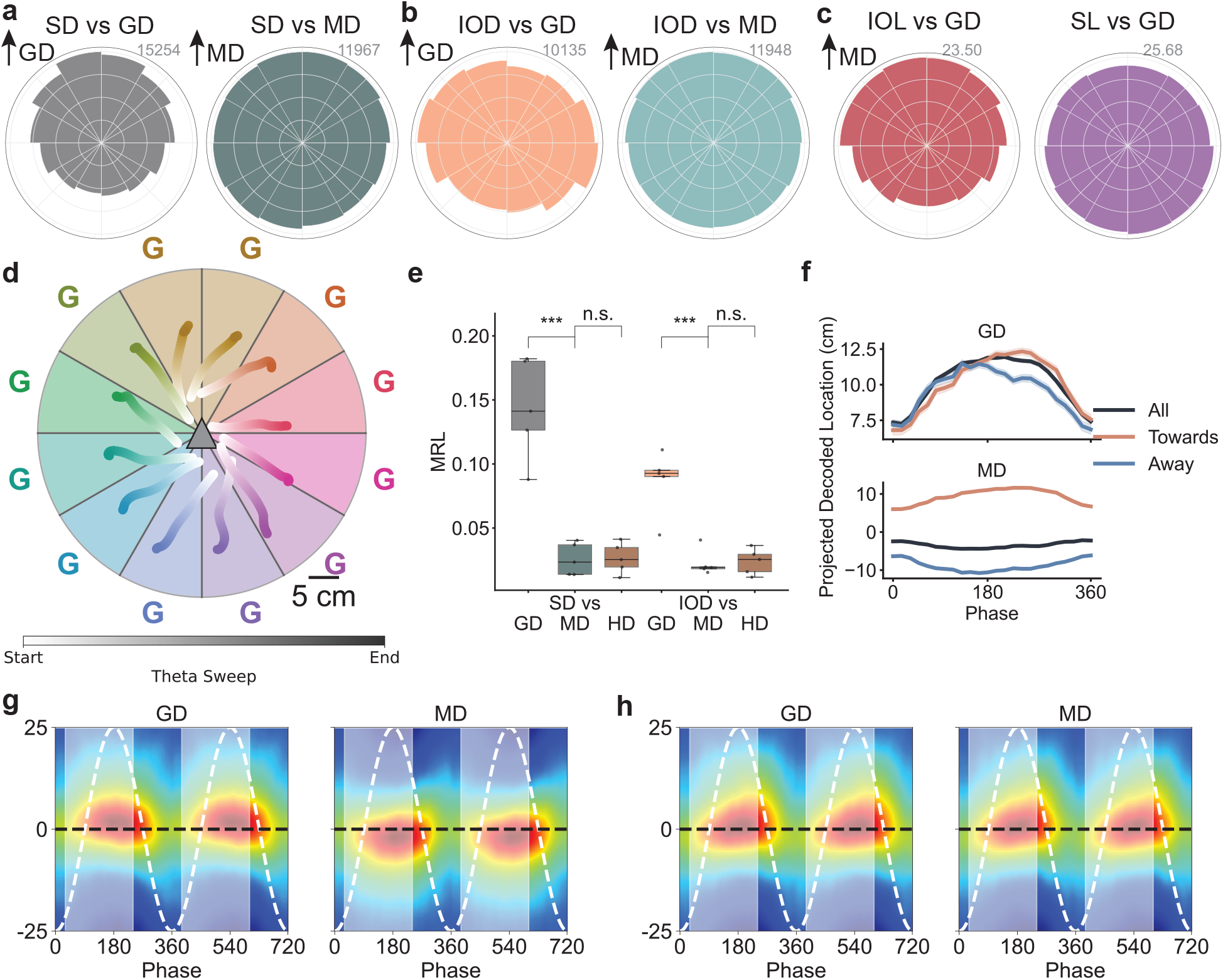
Goal-oriented directional bias in theta sweeps given theta cycles retrieved from multi-unit activity. Theta cycles are derived from LFP in the main analyses. Here we show goal-oriented directional bias persists in theta sweeps given theta cycles retrieved from multi-unit activities (MUA). **a**. Circular histograms of relative angles between SD and GD (left), and SD and MD (right). **b**. Same as (a), but for the IOD. **c**. Median distance from initial offset of decoded theta sweeps to rat’s location (IOL; left), and median length of theta sweeps (right), as a function of movement direction relative to goal direction (MD upwards). **d**. Median theta sweeps as a function of varying relative MD to the goal (MD upwards). **e**. Comparison of rat-specific MRL of the circular distributions in (a) and (b) (one-sided paired t-test SD wrt GD vs MD: p = 0.0001; two-sided paired t-test SD wrt MD vs HD: p = 0.5393; IOD wrt GD vs MD: p = 0.0004; IOD wrt MD vs HD: p = 0.5713). **f**. Mean projection of decoded theta sweeps (± s.e.) along the goal- (left) and movement-directions (right), over all periods and movements towards and away from the goal. We identify robust goal-oriented directional modulation in theta sweeps. **g-h**. Averaged decoding posteriors over all theta sweeps, projected onto corresponding goal- (left) and movement-directions (right), as a function of theta phase, relative to rats’ position (black dashed line) when rats are moving towards (g) and away from (h) the goal. Two theta cycles (white dashed line) are shown. Shaded areas indicate average (over all rats) phase range of “forward-extending” theta sweeps (*9, 24*). Analysis given either MUA- or LFP-derived theta cycles lead to the same main findings of robust goal-oriented directional modulation in theta sweeps.

**Figure S10.**
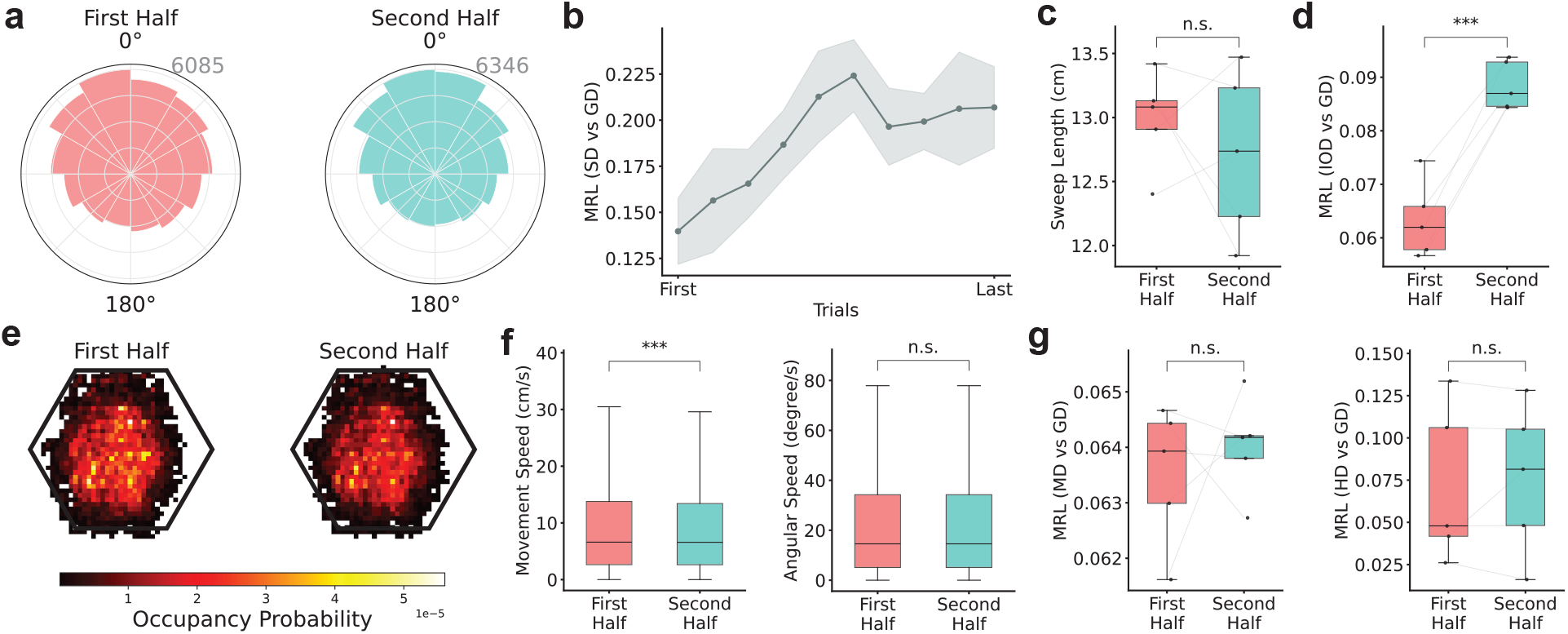
Goal-modulation in theta sweeps becomes stronger with experience. **a**. Circular histograms of relative angles between SD and GD over the first (left) and second (right) half of all trials (aggregating over all rats). **b**. MRL of the circular distribution of relative angles between SD and GD, as a function of trial number (mean MRL over 5 rats, shaded area indicates 1 s.d.). **c**. Rat-specific median length of theta sweeps over the first and second half of all trials (paired two-sided t-test: p = 0.2342). **d**. Rat-specific MRLs of relative angles between IOD and GD over the first and second half of all trials (paired two-sided t-test: p = 0.0005). **e**. Spatial occupancy over the first (left) and second (right) half of all trials (aggregated across all rats). The state occupancy distributions over early and late trials are highly similar (Bhattacharyya coefficient, *ρ* = 0.9909). **f**. Movement (left) and rotational (right) speed of rats over the first and second half of all trials (aggregated across all rats). Rats move significantly faster during early trials comparing to late trials (one-sided t-test: p = 6.8480 × 10^−5^), and there is no significant difference between head rotational speed (two-sided t-test: p = 0.9255). Hence the increased concentration of SD relative to GD is not due to speed confound (c.f. Figure 2b). **g**. Rat-specific MRLs of relative angles between MD and GD (left), and HD and GD (right),over the first and second half of all trials. No significant difference between the spread of the two circular distributions is observed (two-sided paired t-test; MD vs GD: p = 0.9030; HD vs GD: p = 0.3955).

**Figure S11.**
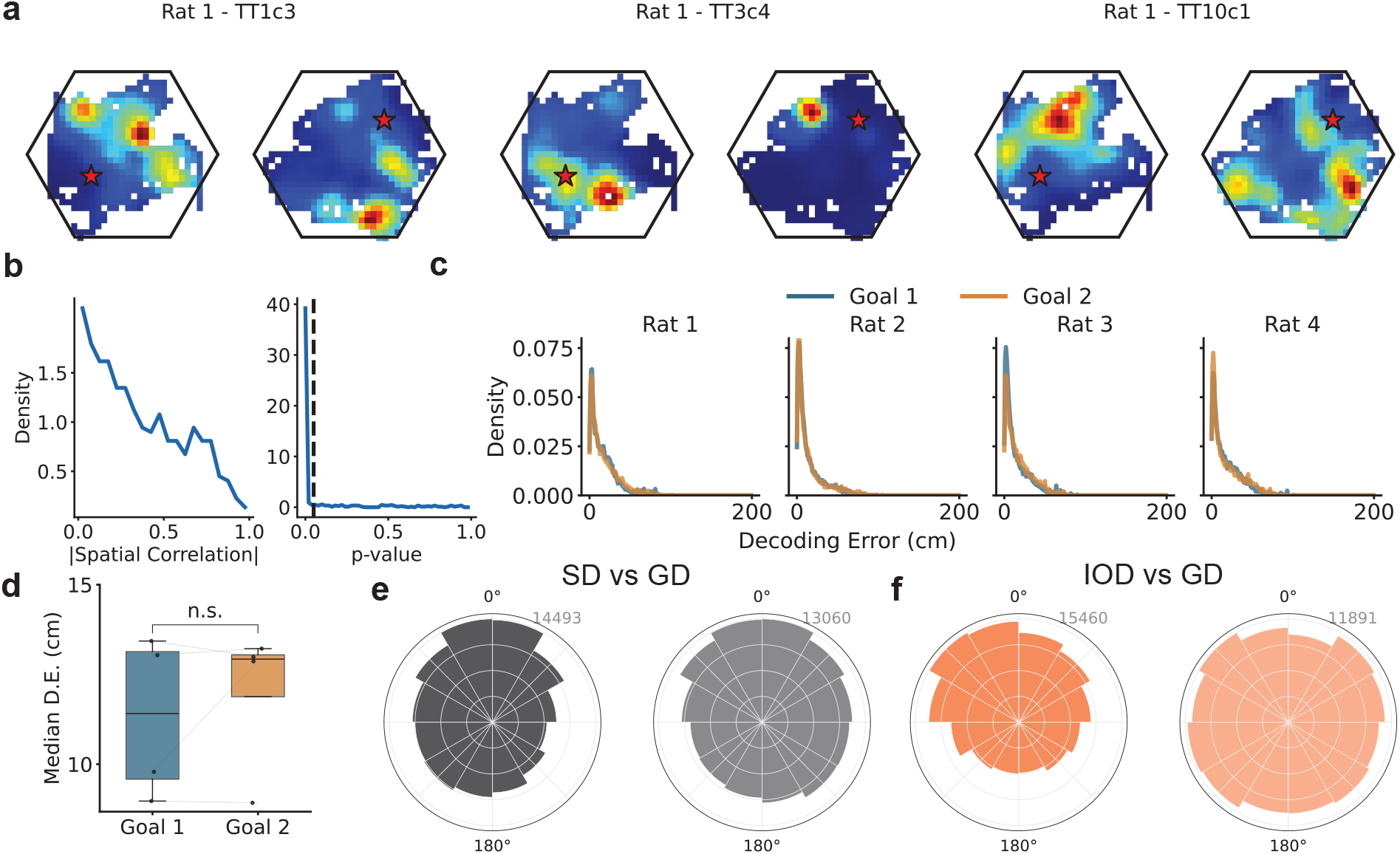
Goal-modulation in theta sweeps is tethered to current goals after goal-switching. **a**. Three example neurons exhibiting remapping corresponding to switching in goal location (left: before; right: after; red star indicates goal locations). **b**. Probability densities of spatial correlations between ratemaps before and after goal-switching (left) and corresponding p-values given permutation tests against chance-level (right), aggregating all cells from Rats 1-3. There are 359/445 (80.7%) cells exhibiting significant changes in spatial tuning (“remapping”) after goal-switching (p < 0.05; permutation test). **c**. Probability densities of decoding errors for Rats 1-4 (from left to right) before (Goal 1) and after (Goal 2) goal-switching. **d**. Comparison of rat-specific median decoding error (DE) in Goal-1 and Goal-2 trials (two-sided paired t-test: p = 0.7733). **e**. Circular histogram of relative angles between SD, and directions to the current (left) and opposing (right) goal locations (aggregating over all trial before and after goal-switching and over all rats). **f**. Same as (e), but for the relative angles with respect to the direction of initial offset of theta sweeps. Theta sweeps extend towards new goal locations post goal-switching, and the initial offset is more sensitive to changes in the goal location.

**Figure S12.**
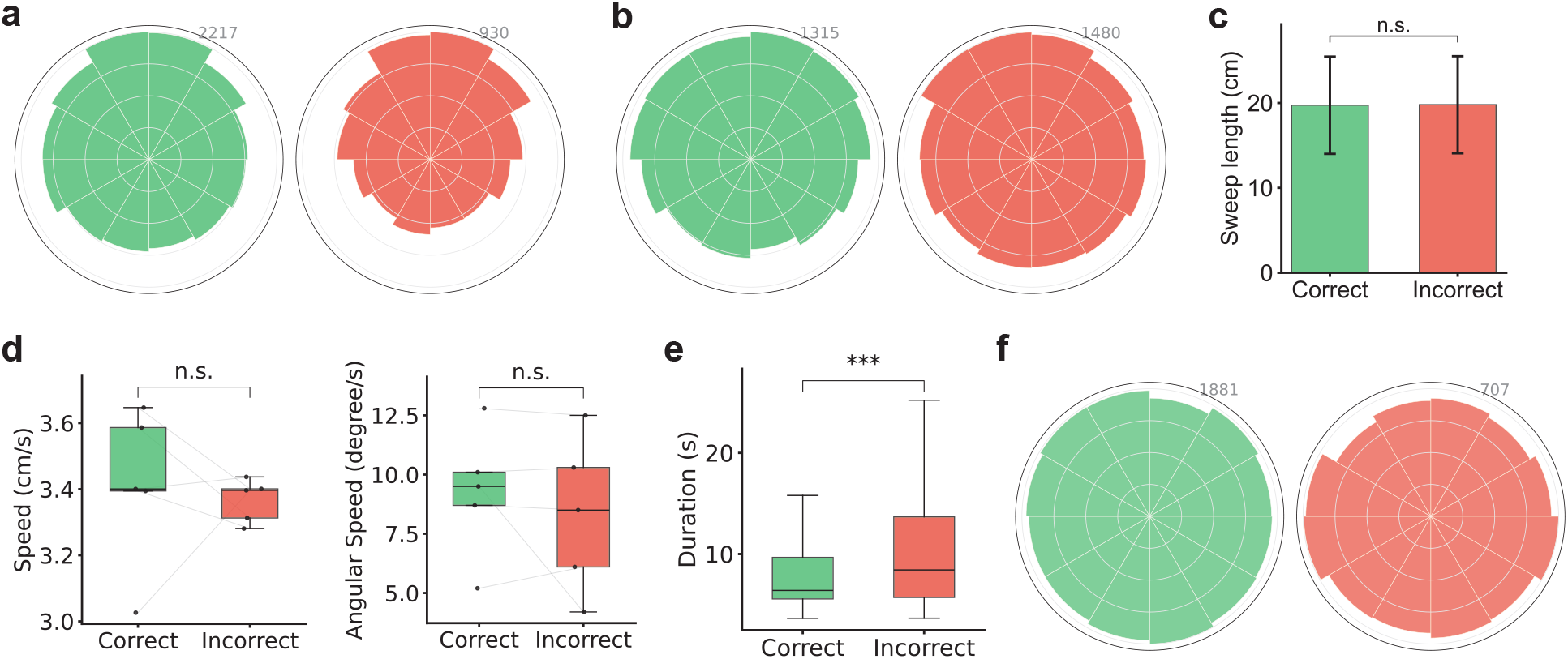
Stronger goal-modulation preceding correct choices. **a**. Circular histograms of relative angles between SD and GD (raw data) during “wait-periods” preceding correct (left) and incorrect (right) choices. **b**. Circular histograms of residual relative angles between SD and GD, after regressing out the effect of distance to the goal (*10, 25*), over correct (left) and incorrect (right) choices. Rat-specific MRL of the distributions is reported in Figure 2f. **c**. Distributions of sweep lengths over correct and incorrect choices (one-sided paired t-test: p = 0.2534). **d**. Movement (left) and head-rotational (right) speed during correct and incorrect choices (two-sided paired t-test; movement speed: p = 0.3596; angular speed: p = 0.2224). **e**. Duration of choice periods over correct and incorrect choices (two-sided paired t-test: p = 0.0002). Rats tend to spend more time before making erroneous decisions, but the frequency of sweep occurrence is similar between correct and incorrect choices (7.6751 sweeps/s and 7.5858 sweeps/s, respectively). **f**. Circular histograms of relative angles between MD and GD over correct (left) and incorrect (right) choices. Apart from distance to the goal, there are no additional significant behavioural differences between correct and incorrect choices that may act as potential confounding factors.

**Figure S13.**
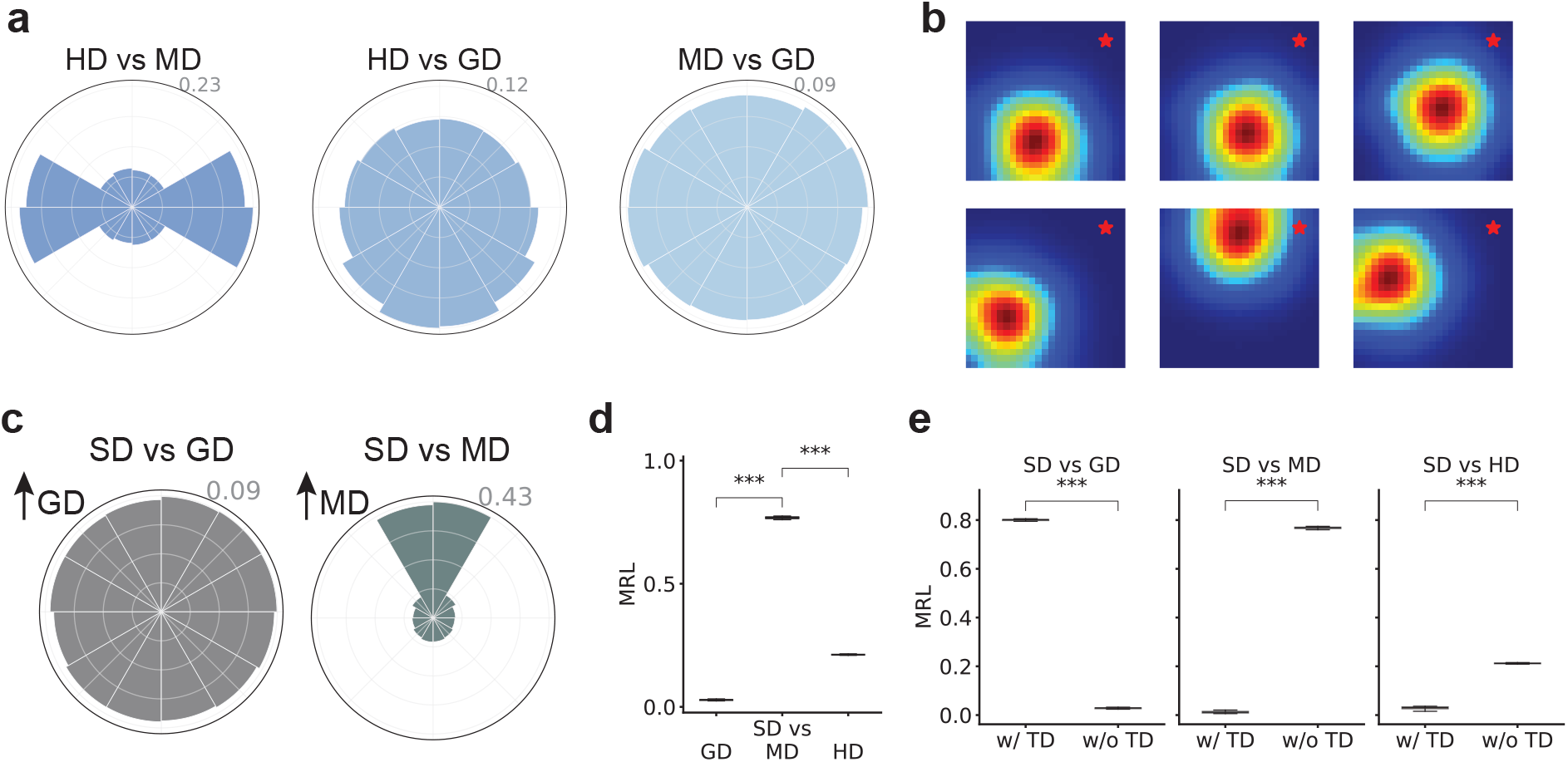
Absence of goal-directed theta sweeps and after turning off the top-down goal-oriented directional input. **a**. Circular histograms of relative angles between HD, MD, and GD over simulated trajectories (Figure 3c). Same behavioural trajectories are used in both simulations with and without top-down inputs. The frequent occurrence of orthogonal HD and MD is due to the simulated rotational scanning (where movement is always perfectly sideways to the heading). Notably, same as the real data in the honeycomb maze, neither headings nor movement exhibits goalward biases. **b**. We perform ablation studies on the effect of goal-oriented directional input. Turning off the top-down input will effectively yield the alternating forward and backward sweeps in the DC network along the internal directions, instead of the goal direction, resulting in left-right alternating sweeps along the movement direction in the activity bump in the downstream PC network (*9, 18*). Example spatial tuning of selected neurons in the PC network exhibits similar unimodal spatial selectivity when the top-down goal-oriented directional input is turned off. **c**. Circular histograms of relative SD with respect to movement- (left) and goal-directions (right). **d**. Comparison of MRLs of the circular distributions of SD vs GD, SD vs MD, and SD vs HD (over 10 random seeds; two-sided paired t-test; SD vs GD: p = 1.1089 × 10^−36^; MD vs HD: p = 1.3159 × 10^−35^). **e**. Comparison of MRLs of the circular distributions of SD vs GD (left), SD vs MD (middle) and SD vs HD (right), when the top-down input is turned on and off, respectively (two-sided paired t-test; SD vs GD: p = 1.9803 × 10^−38^; SD vs MD: p = 3.6838 × 10^−36^; SD vs HD: p = 3.9300 × 10^−25^). As predicted, the top-down inputs is the essential mechanism underlying goal-directed theta sweeps in our simulations, and simulated theta sweeps predominantly following movement direction and exhibit little goal-modulation in its absence.

**Table S1.** Model parameters used in continuous attractor network simulations.

| Component | Parameter | Value | Description |
| --- | --- | --- | --- |
| **Directional Cells Network** | $\tau_h$ | 10 ms | Time constant |
| | $N_h$ | 100 | Number of neurons |
| | $A_h$ | 3.0 | Sensory input strength |
| | $\sigma_h$ | 0.4 radians | Sensory input tuning width |
| | $\bar{\alpha}_h$ | 0.4 | Theta modulation strength |
| | $A_h^{td}$ | 4.0 | Top-down goal-modulated input strength |
| | $\gamma_h$ | 0.4 radians | Top-down goal-modulated input tuning width |
| | $J_0^h$ | 0.8 | Recurrent connection strength |
| | $\eta_h$ | 0.4 radians | Recurrent connection width |
| | $\tau_a^h$ | 100 ms | Adaptation time constant |
| | $m_h$ | 1.0 | Adaptation strength |
| | $k_h$ | 1.0 | Global inhibition strength |
| **Place Cells Network** | $\tau_p$ | 10 ms | Time constant |
| | $N_p$ | 50 | Number of neurons |
| | $\sigma_p$ | 50 cm | Spatial input width |
| | $\bar{\alpha}_p$ | 0.8 | Theta modulation strength |
| | $k_p$ | 1.0 | Global inhibition strength |
| | $g$ | 1000 | Population gain factor |
| | $J_0^p$ | 0.001 | Recurrent connection strength |
| | $\eta_p$ | 50 cm | Recurrent connection width |
| | $\tau_p^a$ | 100 ms | Adaptation time constant |
| | $m_p$ | 0.1 | Adaptation strength |
| | $\gamma_p$ | 50 cm | Conjunctive grid cell input tuning width |
| | $\beta_0$ | 0.2 | Speed scaling factor |
| | $C_p$ | 0.4 | Speed scaling baseline |
| **Place Cells Network (continued)** | $w_0$ | 45 cm | Internal direction shift |
| | $w_1$ | 150 cm | Head direction shift |
| | $\nu$ | 0.99 | Activity threshold |
| **Simulation Parameters** | $dt$ | 1 ms | Time step for numerical integration |
| | $f_\theta$ | 10 Hz | Theta frequency |
|  | *Duration* | 300 s | Simulation duration |

